# A frame and a hotspot in cochlear mechanics

**DOI:** 10.1101/2023.06.29.547111

**Authors:** C. Elliott Strimbu, Lauren A. Chiriboga, Brian L. Frost, Elizabeth S. Olson

## Abstract

Auditory sensation is based in nanoscale vibration of the sensory tissue of the cochlea, the organ of Corti complex (OCC). Motion within the OCC is now observable due to optical coherence tomography. In the cochlear base, in response to sound stimulation, the region that includes the electro-motile outer hair cells (OHC) was observed to move with larger amplitude than the basilar membrane (BM) and surrounding regions. The intense motion is based in active cell mechanics, and the region was termed the “hotspot” (Cooper et al., 2018, Nature comm). In addition to this quantitative distinction, the hotspot moved qualitatively differently than the BM, in that its motion scaled nonlinearly with stimulus level at all frequencies, evincing sub-BF activity. Sub-BF activity enhances non-BF motion; thus the frequency tuning of the hotspot was reduced relative to the BM. Regions that did not exhibit sub-BF activity are here defined as the OCC “frame”. By this definition the frame includes the BM, the medial and lateral OCC, and most significantly, the reticular lamina (RL). The frame concept groups the majority OCC as a structure that is largely shielded from sub-BF activity. This shielding, and how it is achieved, are key to the active frequency tuning of the cochlea. The observation that the RL does not move actively sub-BF indicates that hair cell stereocilia are not exposed to sub-BF activity. A complex difference analysis reveals the motion of the hotspot relative to the frame.

## Introduction

Phase-sensitive optical coherence tomography (OCT) has allowed observations of motion within the sensory tissue of the organ of Corti complex (OCC). Before OCT, observations were restricted to the first-encountered surface of the organ, which is the basilar membrane (BM) in the basal, high-frequency, region discussed in this report. Decades of observations revealed and confirmed that in healthy cochleae, frequency tuning at an auditory neuron is much like that of BM motion (Narayan et al., 1998). BM motion is nonlinear (it scales compressively with sound level) likely due to active outer-hair-cell (OHC)-based forces driven by mechano-electric transduction (MET) current. The MET current saturates, leading to nonlinear OHC-based force (Fettiplace and Kim, 2014; Iwasa and Adachi, 1997). Thus, the non-linear motion is “active” motion. At the BM, activity is limited to frequencies near the best frequency (BF) of the measurement location, where *post-mortem* motion is reduced by factors approaching 1000 at low sound pressure levels (SPL). (BF is defined as the frequency where motion peaked at the lowest SPL where the peak could be identified.) Sub-BF BM motion is unchanged *post-mortem*, and considered “passive” (Rhode, 2007). The OHC electro- mechanical transduction that is presumed responsible for the in vivo activity is not tuned when measured in isolated OHCs (Frank et al., 1999) and the basis for the frequency tuning of nonlinear, active motion in *vivo* is not known, although creative cochlear models can predict this tuning (for example, Nankali et al., 2020; Yoon et al., 2011).

OCT-based measurements revealed motions in the OHC region of the sensory tissue that were greater than BM motion (at all frequencies at low-moderate SPL, and sub-BF at high SPL), leading to the term “hotspot” for the region (Cooper et al., 2018). The aptness of the term is clear in the motion maps from a healthy gerbil cochlea, as seen in Fig.1C-F and especially Fig1.C-E. Multitone stimuli were used, and the response maps are shown at 40 dB SPL (C and D) and 67 dB SPL (E and F), at the BF (D and F) and BF/2 (C and E). As will be shown in later figures, motion within the hotspot also differed from the BM by showing active, nonlinear motion at sub-BF frequencies (Cooper et al., 2018). Sub-BF nonlinearity appeared at relatively low SPL when multitone stimuli were used, likely because the simultaneous presence of near-BF tones led to saturation of MET current at low-moderate SPL. The presence of sub-BF activity can also be perceived when sub-BF responses are reduced *post-mortem* (Strimbu et al., 2020; He et al., 2018; Cho and Puria, 2022). OHC-generated current is nonlinear (both at BF and sub-BF) at moderate SPL with multitone stimuli and is nonlinear sub-BF at high SPL with single-tone stimuli, reaffirming the expectation that nonlinearity in motion responses is due to MET current saturation (Fallah et al., 2019, Dong and Olson, 2013, 2016).

**Figure 1.**
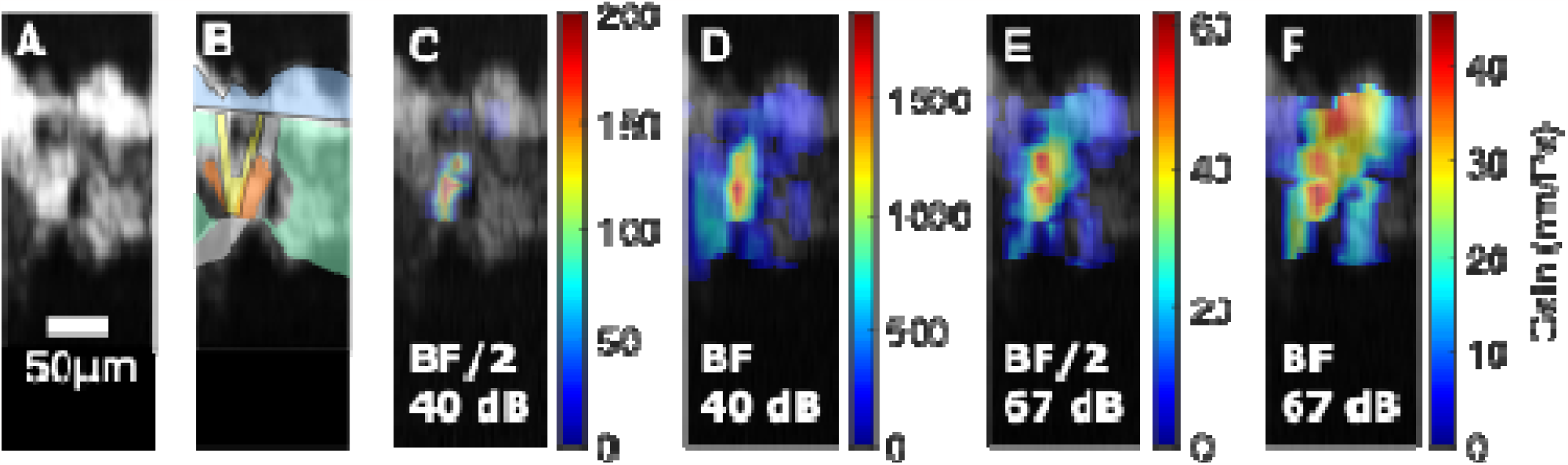
A. Brightness-scan (B-scan). The white regions are regions that reflect the OCT light, the dark regions are fluid spaces. B. B-scan with transparent cartoon of approximate anatomy with blue the BM, yellow the pillar cells, orange the outer and inner hair cells (OHC/IHC to the right/left of pillar cells). The white transparent regions below and above the OHCs are the tectorial membrane (TM) and Deiters cells respectively. (See Fig. 2A for detailed labelling of a radial-transverse cross section.) C-D. Motion response heatmaps for 40 dB SPL multitone stimuli, at the frequencies BF/2 (C) and BF (D). E-F. Motion response heatmaps for 67 dB SPL multitone stimuli, at the frequencies BF/2 (E) and BF (F). Gerbil 995 run 23, BF ~ 31 kHz.

With this background we introduce the concept of the OCC “frame”. The frame is defined as including all locations within the OCC that do not possess sub-BF activity. It is identified in this report as locations that do not exhibit nonlinear sub-BF motion in response to multitone stimuli. The results will show that the frame includes much of the OCC, including regions within the reticular lamina (RL), at the TM-facing surface of the OHCs (Fig. 2A), and regions both lateral and medial to the OHCs. With this observation, stereocilia, which emerge from the RL, are likely not stimulated by sub-BF activity. The side of the OHCs that abuts Deiters cells is termed the OHC\DC region. The discussion analyzes the measured OHC\DC-region motion as a sum of frame and internal-OHC\DC motion and reflects on the ramifications of the presence and absence of active sub-BF motion in different OCC regions.

**Figure 2.**
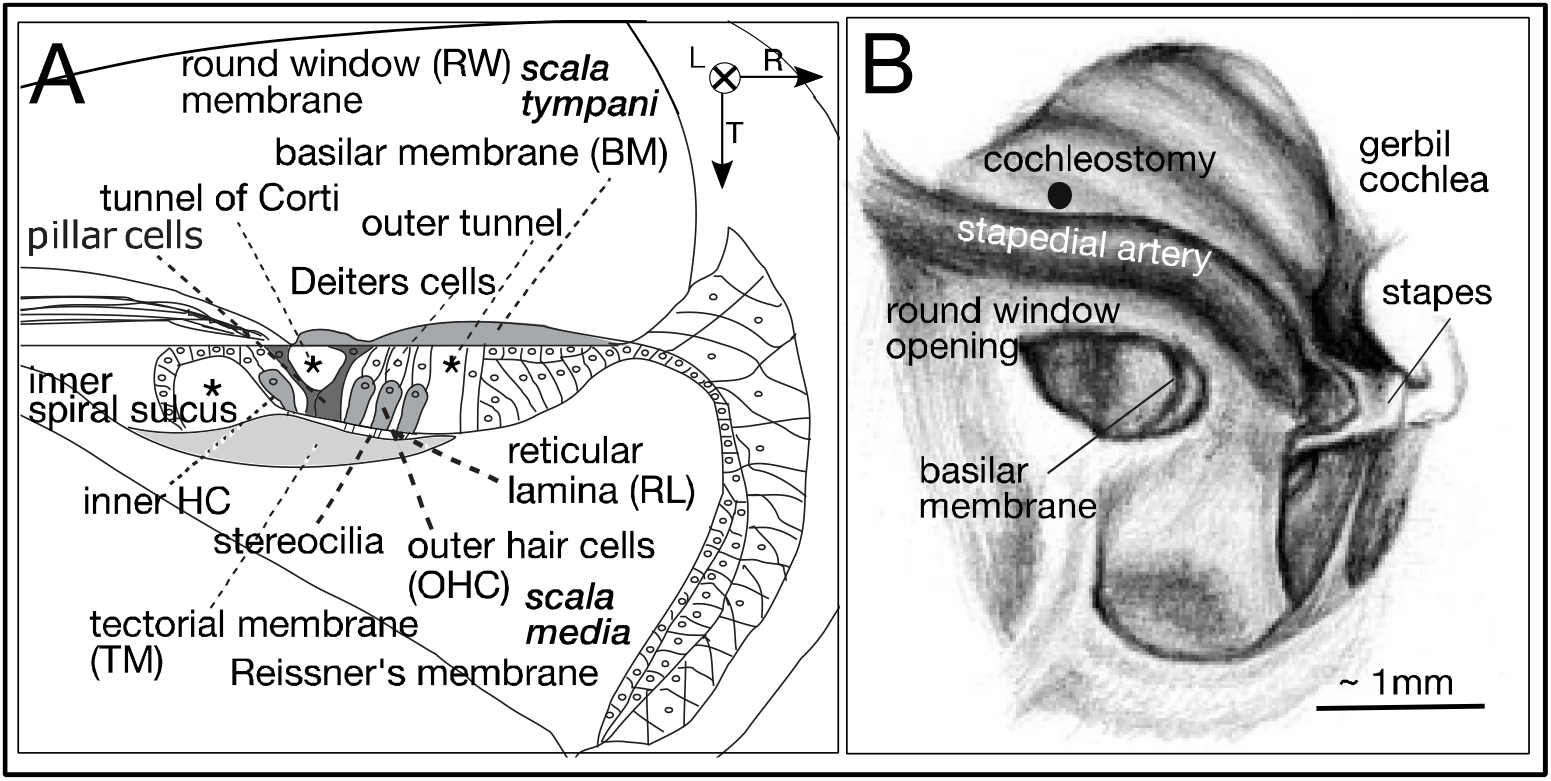
A. Cartoon of a cross section of the gerbil cochlear base; a transverse-radial B-scan gives this view. Longitudinal, radial and transverse directions are defined (L,R,T). The longitudinal direction points along the spiral of the cochlea, from the base towards the apex. Asterisks note the fluid tunnels that appear as dark areas and help parse the anatomy in a B-scan. B. Left gerbil cochlea. Motion measurements were made through the RW membrane, either with the optical axis transverse, in the very basal region, or with the optical axis pointing down the cochlear spiral to reach a lower BF region, with the optical axis including significant longitudinal, transverse and sometimes radial components. The placement of a cochleostomy is shown.

## Methods

The experiments were approved by the Institutional Animal Care and Use Committee of Columbia University. This report includes findings spanning several years of investigation in our lab and more details of the methods can be found in Strimbu et al., 2020.

### Animal preparation

Adult gerbils of both sexes were used. They were initially anesthetized with intraperitoneal (IP) injections of 40 mg/kg ketamine and 40 mg/kg sodium pentobarbital. Buprenorphine, 0.1 mg/kg, was administered at the start of the surgery and after 6 – 8 hours. Supplemental doses of pentobarbital were administered to maintain areflexia in response to a toe pinch. Some animals were also given subcutaneous injections of 2% lidocaine. The animals were tracheotomized to facilitate breathing and their temperatures were maintained with a heating blanket. Supplemental heating was provided by a lamp and disposable hand warmers. The head was attached to a goniometer and the pinna and most of the cartilaginous ear canal (EC) and tissue covering the temporal bones were removed. The bulla was gently opened with forceps. At the end of the experiment the animals were euthanized with pentobarbital. Animal numbering is for internal record keeping.

### Stimulation and response recording

Acoustic stimuli were generated by a Tucker Davis Technologies system and were presented closed-field to the EC by a Radio Shack dynamic speaker. A Sokolich ultrasonic microphone (WGS & Associates, Newport Beach, CA) was coupled to the speaker tube for sound measurement just inside the EC. Measurements were recorded at 98 kHz (97656.25 Hz) or 130 kHz (130208.33 Hz). The multitone stimuli used here were zwuis tone complexes in which *N* frequencies are played simultaneously (van der Heijden and Joris, 2003). The frequencies were chosen to have an integer number of cycles in the recording window, approximately equal spacing (up to a few percent), and had no harmonics or intermodulation distortion products up to third order. Each sinusoidal component of the complex was assigned a random phase so the total sound pressure level, in dB SPL, was ~ 10 log *N* higher than each individual frequency component. The number of frequencies and frequency ranges are indicated in the figure legends. Single-tone stimuli were presented as discreet sweeps in which each frequency was presented for 63 ms including a 1 ms rise/fall tapered with a cosine-squared envelope. Single-tone stimuli were played with a higher maximal level to cover the same dynamic range as the multitone stimuli. Multitone stimuli were 1 s in length except for the data in Fig. 9, which was 10 s. Single-tone sweeps were 2 s total in length.

### DPOAE

As a monitor of cochlear condition, 2f1-f2 distortion product otoacoustic emissions (DPOAEs) were measured at the beginning of the experiment and at several time points during the experiment. The DPOAEs were evoked by two simultaneously presented tones with a fixed ratio f2/f1 = 1.2 and equal levels of 50 and 70 dB SPL. The signals played for 1 second, responses were divided into 50 identical presentations and averaged.

### OCT

Displacement measurements were acquired using a Thorlabs Telesto 320 spectral domain OCT system with a central wavelength of 1300 nm. OCT produces a 1-D “image” within the depth of the preparation, which is termed an axial scan (A-scan); scanning with mirrors along a second axis produces a 2-D image termed a B-scan. Moving the scanning mirrors in a third, orthogonal direction, the instrument can construct a 3-dimensional volumetric scan. The ThorImage program was used to generate B-scans to orient the preparation and locate the regions of interest within the cochlea, and for volumetric (3-D) imaging. The OCT system is equipped with an LSM03 objective lens, and the properties of the light source and lens determine the system’s imaging resolution. The system has an axial resolution of approximately 4 μm and a lateral resolution of approximately 10 μm. The vertical positions of measurement locations in a B-scan are noted in pixels, and the distance between adjacent pixels is 2.7 μm. A time series of A-scans (Motion-scans or M-scans) was taken at selected regions in the cochlea, to measure vibration. Acquisition and initial processing of the OCT vibrometry data were performed using custom software written in C++ using the Thorlabs Spectral Radar software development kit.

### Experimental design

Measurements made with an M-scan along a single axis were typically made close to the intersection of the arcuate and pectinate zones, passing through the outer hair cell region. B-scan area-spanning recordings of the uniaxial vibrations were constructed by taking sequential M-scans in 10 μm quasi-radial steps, termed “slices”. To confirm that the structures were stable over the length of the recordings, the OCT software took B-scans of the same region before and after each set of vibration measurements and we compared the two images to exclude recordings with significant drift. Further offline analysis was performed in custom software written in MATLAB. Once the vibration time waveforms were acquired for the regions of interest, the amplitudes and phases at the sound stimulus frequencies were extracted by Fourier analysis. For each stimulus frequency, the response was deemed significant if the Fourier coefficient was 3 times larger than the standard deviation of the noise level, measured from 10 neighboring bins in the spectra. Responses are reported as gain relative to ear canal pressure in nm/Pa, and phase relative to ear canal pressure in cycles.

This report includes OCT-observations made over several years in several studies. None of the specific data here were previously published. A common aspect of experimental design was to simultaneously observe motion at different locations within the sensory tissue of the OCC. The studies sometimes pursued specific hypotheses, for example regarding the effect of ototoxic drugs (Strimbu et al. 2020, 2022), but the results here are of unperturbed cochleae.

## Analytical methods

### Longitudinal, radial, transverse components of the optical axis

Frost et al., 2022 outlined a program in which a volume scan is used to generate the mapping between anatomical coordinates (the longitudinal (L), radial (R) and transverse (T) directions noted in Fig. 2A) and optical coordinates from an OCT volume scan. For some of the data sets here L, R, T components of the optical axis are reported. The relative values of these components indicate how much each of these directions is represented in the measured motion. For example, in Fig. 4, the relative L, R, T components were 0.85,0.24,0.47, signifying that the optical axis was primarily in the longitudinal direction but with substantial radial and transverse contributions. (Two significant figures are excessive for these results but are retained for clarity, so that the components sum in quadrature to ~ 1.) Motion along any of the anatomical axes would contribute to the displacement measurement, with the actual motion in each anatomical direction weighted by its respective signed component in the reported motion.

### Metrics related to quantifying sub-BF activity

Metrics are defined to support the division into “sub-BF active” (hotspot) and “sub-BF not-active” (frame) regions. As will be discussed with respect to Fig. 4, the metrics are applied to data taken with multitone stimuli. The following measured quantities are used:

- The response gain at BF, measured at 70 dB SPL, *G*_*BF70*_
- The response gain at BF/2, measured at 70 dB SPL, *G*_*0*.*5BF70*_ (In gerbil 995 the measurements were done at 67 instead of 70 dB SPL, and 67 dB SPL data are used.)
- The response gain at BF/2, measured at 80 dB SPL, *G*_*0*.*5BF80*_

Our first metric for assessing sub-BF activity is *GR1*:

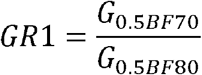

If sub-BF nonlinearity is perfectly absent, *GR1* = 1. If compressive nonlinearity is present, *GR1* > 1. Due to random fluctuations in data, *GR1* never exactly equals 1 and we choose a value of 1.3 as a cutoff, with GR1values greater than 1.3 showing sufficient nonlinearity to be considered sub-BF active. This metric was sometimes ambiguous, leading us to define a second metric, *GR2(dB):*

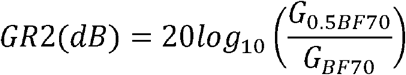

The rationale for *GR2(dB)* and its logarithmic form are as follows: At moderate-high SPL (we use 67 or 70 dB), locations that show sub-BF nonlinearity typically have higher gain at BF/2 than at BF, resulting in ratio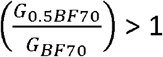. The locations that do not show sub-BF nonlinearity typically have higher gain at BF than BF/2, an 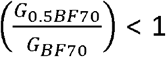. We take the logarithm (presented as dB value) because that neatly divides sub-BF active and sub-BF not-active into positive and negative *cR2(dB)* values respectively. We use these two metrics to support the categorization into sub-BF active locations and sub-BF non-active locations. Table 1 has gain metrics for all the presented data and some additional locations. In a few instances the two metrics were ambiguous regarding whether or not a location was in the frame. This is pointed out below for a few locations.

**Table 1.** Metrics used to quantitatively separate locations in which sub-BF activity was present from locations in which sub-BF activity was absent. The first metric is GR1, the gain at 70 dB SPL divided by the gain at 80 dB SPL, both measured at BF/2. This is a straightforward measure of nonlinearity, and if nonlinearity were absent, GR1 would equal 1 If compressive nonlinearity were present, GR1 would be greater than 1. Due to experimental data fluctuations, GR1 is never exactly equal to 1. We used GR = 1.3 as a cut-off for the presence of sub-BF compressive nonlinearity. Because fluctuations could lead GR1 to equal 1.3 (and even 1.4 in G996 run5 px 94) at BF/2, while adjacent frequencies were more clearly linear, we adopted a second metric to address these ambiguous locations. GR2(dB) is the ratio of the gain at 70 dB at BF/2 to the gain at 70 dB at BF, expressed in dB. At locations with sub-BF compressive nonlinearity, the increased BF/2 gain would tend to make this ratio value greater than 1. By taking the dB value, positive GR2(dB) values correspond to sub-BF activity, and negative GR2(dB) values to sub-BF lack of activity. In experiment 995 we gathered data at 67 instead of 70 dB SPL, thus in this experiment the measurements were done at 67 and 80 dB SPL. (This was done in order to gather data over four SPLs, with the lowest level 40 dB (rather than the usual 50) and the highest 80.) In the table red color indicates locations with sub-BF activity, pink indicates locations that remained ambiguous and gray/white are locations without sub-BF activity.

| ID | Pixel | GR1 | GR2(dB) | ID | Pixel | GR1 | GR2(dB) | ID | Pixel | GR1 | GR2(dB) |
| --- | --- | --- | --- | --- | --- | --- | --- | --- | --- | --- | --- |
| G986 r4 | 51 | 1.0 | -4 | G995 r23 slice 3 | 157 | 1.0 | -6 | G995 r23 slice 9 | 148 | 0.9 | -6 |
| G986 r4 | 53 | 1.2 | -3 | G995 r23 slice 3 | 174 | 2.1 | 0 | G995 r23 slice 9 | 153 | 0.8 | -6 |
| G986 r4 | 55 | 1.3 | -3 | G995 r23 slice 3 | 178 | 2.5 | 1 | G995 r23 slice 9 | 160 | 0.8 | -6 |
| G986 r4 | 60 | 0.8 | -7 | G995 r23 slice 3 | 186 | 1.8 | -4 | G995 r23 slice 9 | 166 | 0.9 | -5 |
| G986 r4 | 65 | 1.9 | 11 | G995 r23 slice 3 | 197 | 1.6 | -2 | G995 r23 slice 9 | 185 | 0.7 | -5 |
| G986 r4 | 69 | 1.5 | 13 | G995 r23 slice 4 | 153 | 0.7 | -3 | G995 r23 slice 9 | 196 | 1.0 | -1 |
| G986 r4 | 74 | 1.3 | 10 | G995 r23 slice 4 | 156 | 0.7 | -4 | G995 r23 slice 10 | 150 | 0.9 | -6 |
| G986 r4 | 76 | 0.7 | -16 | G995 r23 slice 4 | 159 | 0.9 | -2 | G995 r23 slice 10 | 153 | 0.8 | -6 |
| G963 r5 | 172 | 1.0 | -10 | G995 r23 slice 4 | 177 | 1.6 | 3 | G995 r23 slice 10 | 161 | 0.8 | -6 |
| G963 r5 | 184 | 1.5 | 4 | G995 r23 slice 4 | 184 | 1.8 | 3 | G995 r23 slice 10 | 163 | 0.8 | -6 |
| G963 r5 | 190 | 0.7 | -1 | G995 r23 slice 4 | 189 | 1.8 | 1 | G995 r23 slice 10 | 201 | 0.7 | -2 |
| G963 r5 | 192 | 0.9 | -24 | G995 r23 slice 4 | 191 | 1.3 | -2 | G995 r23 slice 11 | 153 | 0.8 | -6 |
| G959 r12 | 122 | 0.8 | -6 | G995 r23 slice 5 | 157 | 0.8 | -1 | G995 r23 slice 11 | 155 | 0.7 | -7 |
| G959 r12 | 133 | 0.8 | -8 | G995 r23 slice 5 | 159 | 0.8 | -7 | G995 r23 slice 11 | 161 | 0.9 | -5 |
| G959 r12 | 138 | 0.9 | -7 | G995 r23 slice 5 | 164 | 1.0 | -4 | G995 r23 slice 11 | 165 | 0.9 | -5 |
| G959 r12 | 146 | 0.9 | -6 | G995 r23 slice 5 | 166 | 0.8 | -6 | G995 r23 slice 12 | 162 | 0.8 | -8 |
| G959 r32 | 100 | 0.9 | -5 | G995 r23 slice 5 | 171 | 2.5 | 4 |  |  |  |  |
| G959 r32 | 107 | 0.9 | -6 | G995 r23 slice 5 | 174 | 2.1 | 2 | G996 r8 slice 5 | 88 | 0.9 | -14 |
| G959 r32 | 115 | 1.6 | 9 | G995 r23 slice 5 | 177 | 1.6 | 2 | G996 r8 slice 5 | 94 | 1.4 | -10 |
| G959 r32 | 122 | 1.6 | 16 | G995 r23 slice 5 | 180 | 2.4 | 2 | G996 r8 slice 6 | 86 | 1.1 | -12 |
| G959 r32 | 128 | 1.6 | 12 | G995 r23 slice 5 | 185 | 2.9 | 0 | G996 r8 slice 6 | 90 | 1.1 | -11 |
| G959 r32 | 144 | no BF/2 data see text |  | G995 r23 slice 5 | 187 | 2.8 | 1 | G996 r8 slice 6 | 94 | 1.0 | -9 |
| G750 r5 | 225 | 0.9 | -4 | G995 r23 slice 6 | 157 | 0.8 | -6 | G996 r8 slice 6 | 103 | 1.2 | -16 |
| G750 r5 | 230 | 0.9 | -3 | G995 r23 slice 6 | 162 | 1.0 | -4 | G996 r8 slice 7 | 86 | 1.1 | -11 |
| G750 r5 | 233 | 0.9 | -3 | G995 r23 slice 6 | 168 | 2.0 | 3 | G996 r8 slice 7 | 93 | 1.7 | 8 |
| G750 r5 | 242 | 1.7 | 10 | G995 r23 slice 6 | 172 | 2.3 | 3 | G996 r8 slice 7 | 96 | 1.9 | 4 |
| G750 r5 | 247 | 1.6 | 12 | G995 r23 slice 6 | 176 | 2.2 | 3 | G996 r8 slice 7 | 104 | 0.8 | -21 |
| G750 r5 | 250 | 1.7 | 12 | G995 r23 slice 6 | 179 | 2.0 | 3 | G996 r8 slice 8 | 80 | 0.9 | -14 |
| G750 r5 | 255 | 1.7 | 13 | G995 r23 slice 6 | 183 | 2.2 | 3 | G996 r8 slice 8 | 84 | 1.2 | -11 |
| G750 r5 | 279 | 0.8 | 1 | G995 r23 slice 6 | 185 | 2.1 | 4 | G996 r8 slice 8 | 87 | 1.3 | -9 |
| G750 r5 | 285 | 0.7 | 0 | G995 r23 slice 7 | 153 | 0.9 | -7 | G996 r8 slice 8 | 95 | 2.4 | no 70 dB BF data |
| G750 r12 | 105 | 1.0 | 4 | G995 r23 slice 7 | 155 | 0.9 | -6 | G996 r8 slice 9 | 81 | 1.2 | -12 |
| G750 r12 | 208 | 0.9 | 0 | G995 r23 slice 7 | 159 | 0.8 | -7 | G996 r8 slice 9 | 86 | 1.0 | -12 |
| G750 r12 | 211 | 1.0 | -4 | G995 r23 slice 7 | 161 | 0.8 | -6 | G996 r8 slice 9 | 107 | 2.8 | -8 |
| G750 r12 | 221 | 0.9 | -4 | G995 r23 slice 7 | 172 | 1.4 | 0 | G996 r8 slice 10 | 79 | 1.1 | -11 |
| G750 r12 | 229 | 1.0 | -6 | G995 r23 slice 7 | 200 | 1.2 | -3 | G996 r8 slice 10 | 82 | 1.1 | -12 |
| G750 r12 | 235 | 0.8 | -5 | G995 r23 slice 8 | 151 | 0.9 | -7 | G996 r8 slice 10 | 84 | 1.1 | -11 |
| G750 r12 | 241 | 0.9 | -4 | G995 r23 slice 8 | 154 | 0.8 | -6 | G996 r8 slice 10 | 87 | 1.2 | -12 |
| G750 r12 | 247 | 0.6 | -4 | G995 r23 slice 8 | 159 | 0.8 | -6 | G996 r8 slice 11 | 84 | 1.1 | -11 |
| G750 r12 | 259 | 0.9 | 2 | G995 r23 slice 8 | 163 | 0.9 | -5 | G996 r8 slice 11 | 87 | 1.1 | -12 |
| G750 r12 | 265 | 0.6 | 3 | G995 r23 slice 8 | 173 | 0.9 | -6 | G996 r8 slice 11 | 98 | 1.3 | -6 |
|  |  |  |  | G995 r23 slice 8 | 176 | 1.1 | -1 | G996 r8 slice 12 | 81 | 0.9 | -13 |
| G995 r23 slice 1 | 156 | 1.2 | -4 | G995 r23 slice 8 | 182 | 0.9 | -6 | G996 r8 slice 12 | 85 | 1.0 | -13 |
| G995 r23 slice 1 | 161 | 0.9 | -4 | G995 r23 slice 8 | 184 | 1.4 | -2 | G996 r8 slice 12 | 89 | 1.0 | -13 |
| G995 r23 slice 2 | 159 | 0.8 | -3 | G995 r23 slice 8 | 188 | 0.7 | -1 | G996 r8 slice 13 | 83 | 1.1 | -11 |
| G995 r23 slice 2 | 162 | 1.1 | 0 |  |  |  |  | G996 r8 slice 13 | 86 | 1.1 | -12 |
| G995 r23 slice 2 | 165 | 1.3 | 3 |  |  |  |  | G996 r8 slice 13 | 88 | 0.7 | -13 |
|  |  |  |  |  |  |  |  | G996 r8 slice 13 | 92 | 1.1 | -11 |

## Results

### A. DPOAE results

DPOAE responses were measured after opening the bulla and at several times during an experiment. The responses typically stayed steady over hours of data collection. Initial DPOAE responses are shown in Fig. 3. In experiment 959, a cochleostomy was made prior to the DPOAE measurement, which might have produced the relatively low responses above 30 kHz, and the dip close to 20 kHz. In this experiment intracochlear motion responses were measured in the 20-25 kHz BF region in this cochlea, where the DPOAE levels were nearly within the normal range. The DPOAE responses in Fig. 3 are consistent with response levels in previous studies, for example Dong and Olson (2008).

**Figure 3.**
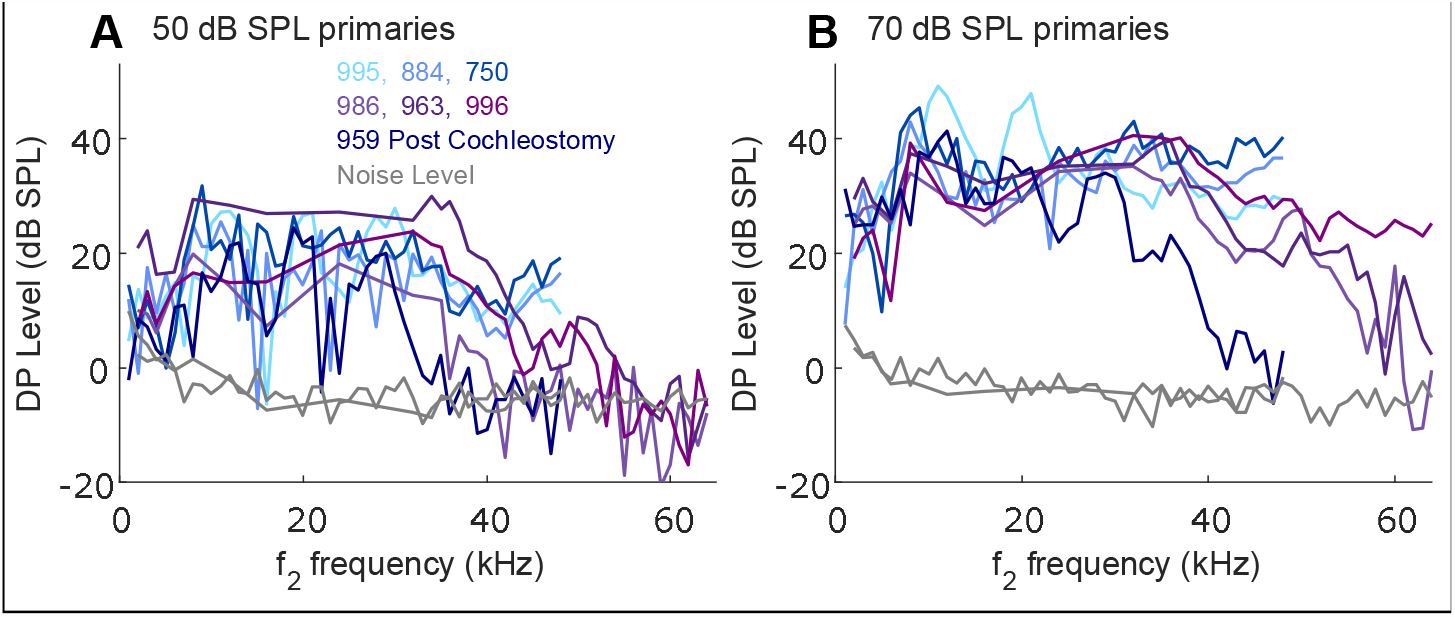
DPOAE responses from the experiments in this report, measured after opening the bulla. A. 50 dB SPL primaries B. **70 dB SPL primaries. The different colors correspond to different experiments, listed in the legend in (A). Blues are** experiments with motion measured at the 25 kHz region and purples from the hook region. One (typical) noise floor is shown for each type of experiment, and the noise levels were very consistent across experiments. The DPOAEs were measured through f2 values of 65 or 45 kHz. A cochleostomy had been made in cochlea 959.

### B. Comparison of multitone and single-tone responses

Fig. 4 illustrates the multitone versus single-tone difference in motion responses. It shows frequency response gains at the BM (Fig. 4A-C) and the OHC\DC region (Fig. 4D-F) with single-tone (Fig. 4A&D) and multitone (Fig. 4B&E) stimuli. Responses are shown to stimuli spanning 45 to 95 dB SPL in the single-tone measurements, and 40 to 80 dB SPL in the multitone measurements. The BF was ~30 kHz; all responses within the BF peak were compressively nonlinear. At the BM, for both single and multitone stimuli, all gains were equal at sub-BF frequencies below 20 kHz. These responses are passive; they are as they would be without activity. At the BM the 95 dB SPL single-tone gains and 80 dB SPL multitone gains were identical through the full frequency range, including the BF (comparison in Fig. 4C). These responses can also be considered passive; in this case the passive responses to sound have dominated active contributions, which have saturated at a level that makes them insignificant (distortion, not discussed here, would still be present).

**Figure 4.**
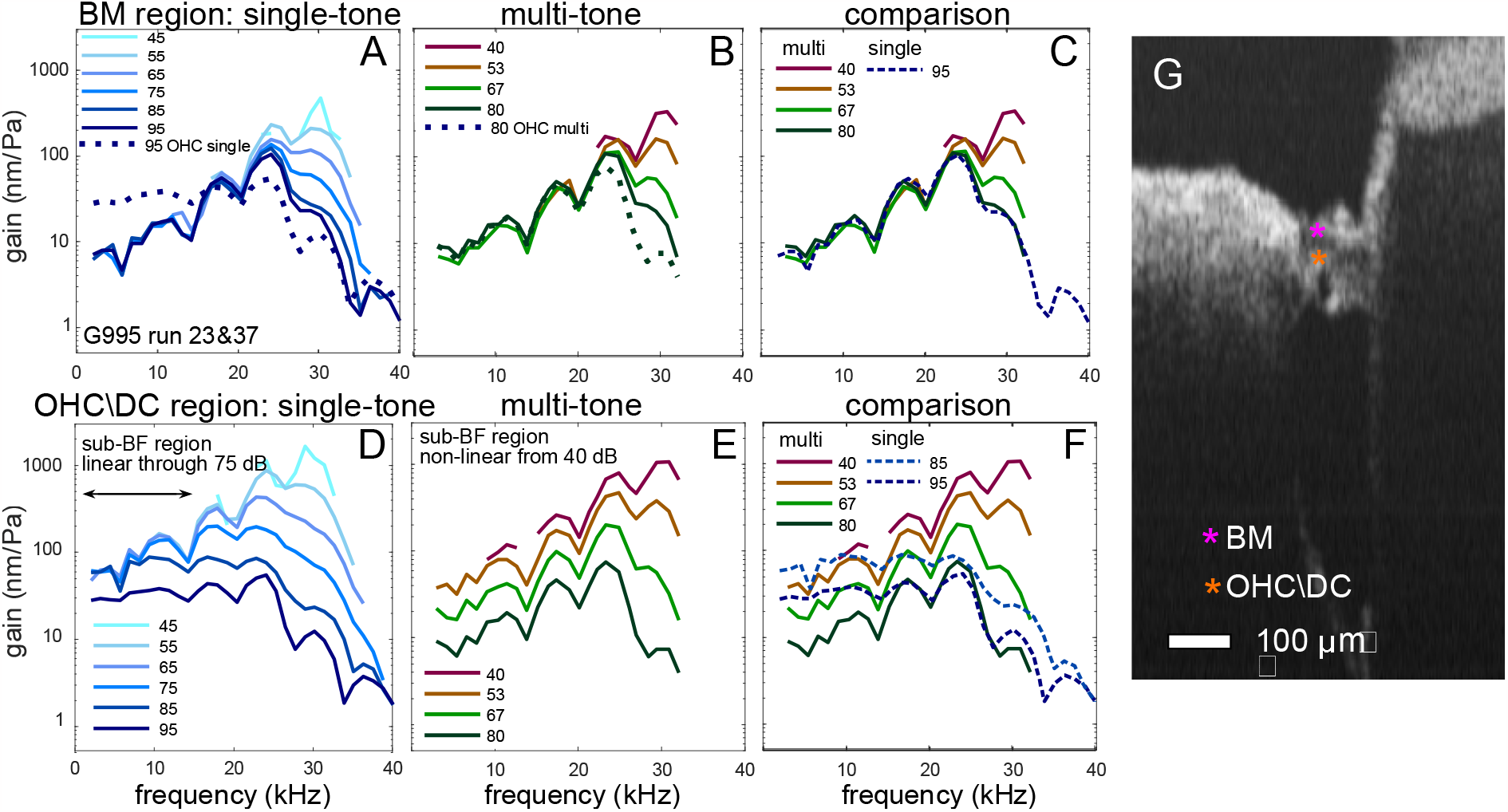
A-C. BM responses to (A) single and (B) multitone stimuli with comparison in (C). D-F. OHC\DC responses to (D) single and (E) multitone stimuli with comparison in (F). G. B-scan showing locations of displayed motion responses. Responses are normalized to EC pressure as gains, so nonlinearity is apparent when curves at different stimulation levels do not overlap. Data were taken through the gerbil RW, with the optical axis aimed apically to reach the ~ 30 kHz BF location. The optical axis had longitudinal, radial and transverse components of 0.85, 0.24, 0.47. The multitone stimuli comprised 25 frequency components, spanning 3 to 32 kHz. Gerbil 995 runs 23&37.

In the OHC\DC-region, the single-tone responses (Fig. 4D) scaled linearly up to 75 dB SPL at sub-BF frequencies below 15 kHz, whereas the multitone responses (Fig. 4E) already showed sub-BF nonlinear scaling at 53 dB SPL. The dashed lines in Fig. 4F show single/multitone comparisons, and the dotted lines in Fig. 4 A&B show OHC\DC to BM comparisons, and support findings from previous studies of BM motion and OHC current. These observations can be summed up as: Multi-tone stimuli are overall higher sound level than their single-tone counterparts and thus produce saturation at lower levels (Fallah et al., 2019; Versteegh et al., 2012). For the purposes of this paper, Fig. 4 D&E demonstrate that multitone are better than single-tone stimuli at producing the sub-BF nonlinearity, which clearly indicates sub-BF activity. In contrast, sub-BF linearity in response to single-tone stimuli, as in the responses up to 75 dB SPL in Fig. 4D, is ambiguous with respect to sub-BF activity. Multitone stimuli were used in what follows.

### C. Results to illustrate the concept of the frame

The remainder of this section consists of examples of motion measurements, chosen to illustrate the concept of the frame. Motion is reported at longitudinal locations directly adjacent to the round window, with a view that is approximately transverse (~45-50 kHz BF place) and at more apical locations, accessed by viewing down the cochlear spiral from the RW, with the optical axis containing a substantial longitudinal component (~24-32 kHz BF place). One data set is included that was taken through a cochleostomy in gerbil, affording a nearly transverse view at the ~ 24 kHz place. The presence of sub-BF activity was evaluated as described in the metrics section of the methods.

### 1. Optical axis in relatively transverse direction

Fig. 5A shows motion gain data sets, Fig. 5B shows the associated A-scan and B-scan. The measurements were made through the RW opening along the dotted line in B. The positions of the measurements spanned 25 pixels, corresponding to 68 μm. Referring to the pixel locations and the A and B-scans in Fig. 5B, pixels 51,53,55 were within the BM, pixels 65 and 69 were in the OHC\DC region and 76 was at the RL-region. Pixel 74 was only 6 μm from pixel 76, and anatomically within the OHC body close to the RL. The sub-BF responses at pixels 65,69 and 74 (OHC\DC) were elevated relative to those at pixels 51-55 (BM), and pixel 76 (RL), and sub-BF nonlinearity (activity) was clear at pixels 65, 69 and 74. Those locations span 9 pixels, ~24 μm. Those locations not exhibiting sub-BF nonlinearity (pixels 51, 53, 55, 60 and 76) were nonlinear near BF and showed a BF peak at 70 dB SPL that become higher as SPL was reduced (when data were available).

**Figure 5.**
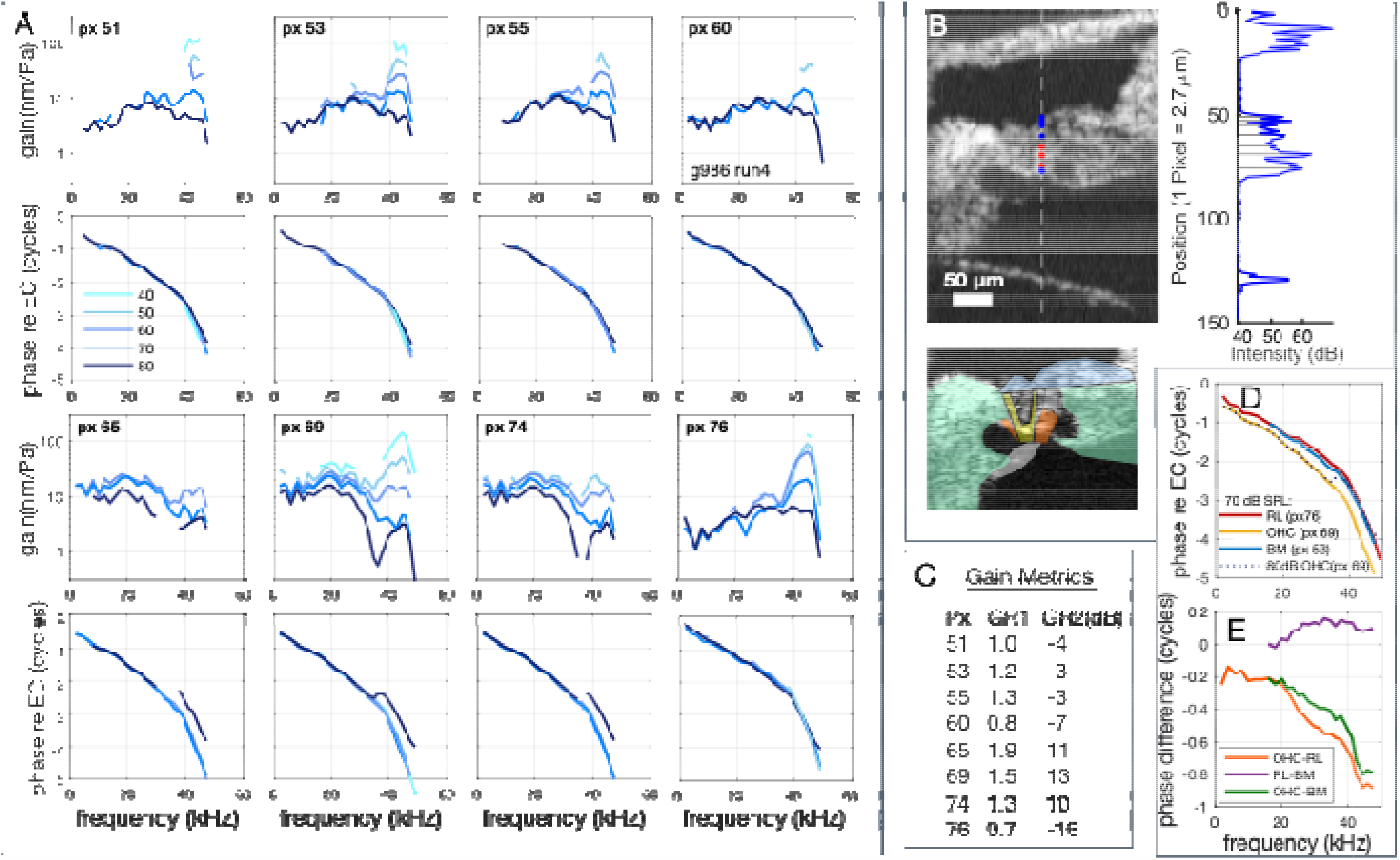
A. Results from eight positions along one axial scan (A-scan). The top panels show gain (displacemen pressure), bottom panels show phase relative to ear canal (EC) pressure. B. B-scan and A-scan for this data set. The da hed line in the B-scan is the displacement measurement axis, with the blue and red dots indicating locations without and with substantial sub-BF nonlinearity/activity. Motion responses were evaluated at locations of peaks in the A-scan (Lin e al., 2017). The A-scan is registered to the B-scan to approximately identify the anatomical locations of measurement. The lower B-scan is anatomically color-coded as in Fig. 1. C. Gain metrics. D. Phases measured at 70 dB SPL from three representative locations. E. Phase differences. The multitone stimuli comprised 35 frequency components, spanning 2 to 60 kHz. The optical axis longitudinal, radial and transverse components were −0.48, −0.35, 0.8. Gerbil 986 run4 BF 44 kHz.

Some oddities of the data are the amplitude trough at high SPL in the OHC\DC region (pixels 65,69,74), occurring along with a phase shift. (The word “trough” is used rather than “notch” because the reduction can span a relatively wide frequency range.) The complex difference analysis in the first part of the discussion will show that this behavior emerges as the result of internal OHC\DC motion riding on the motion of the BM. Another curiosity, addressed here briefly, is that the motions within the OHC (pixels 65, 69, 74) were very similar in magnitude, whereas OHC stretching/compressing would predict a smaller motion close to the RL than further along the OHC body. Experiments done in isolated OHCs in a microchamber help but do not fully resolve this puzzle. They showed that the motion within 10% of the distance to the apical pole of the OHC accounted for ~ 50% of the maximum electromotile motion (Hallworth et al., 1993). Thus, even in isolated OHCs electromotility was not uniformly increasing along the OHC length, and it is not clear what to expect for OHCs coupled with the OCC (Rabbitt, 2022).

Fig. 5D shows the phases from the three different anatomical locations, pixels 53 (BM), 69 (OHC\DC) and 76 (RL), at 70 dB SPL, and phase differences are shown in Fig. 5E. 70 dB SPL data are shown because there was not much variation between SPLs except at 80 dB SPL and the 70 dB SPL data are out of the noise through a relatively wide frequency range. In Fig. 7D, 80 dB SPL OHC\DC region phase is included and shows that at frequencies above the phase shift, the OHC\DC phase was almost in line with the BM and RL phases. The RL and BM phases were similar but not identical, with the RL phase leading the BM phase by a value that increased from ~ 0 to ~ 0.15 cycle from 18 to 35 kHz. The OHC\DC region was close to half a cycle out of phase with the RL (varying between 0.4 and 0.6 cycles difference) from 20 to 40 kHz.

Fig. 5C is a table of metrics as defined in the methods. *GR1*, a ratio of gains at 70 and 80 dB SPL is a direct measure of nonlinearity. *GR1* can be ambiguous due to random data fluctuations. If *GR1*>1.3, sub-BF activity was taken to be supported. If ambiguity needed to be addressed, *GR2(dB)* is used, which compares gains at BF/2 to gains at BF, at 70 dB SPL. Positive *GR2(dB)* values support sub-BF activity. We use the Fig. 5 data set to explore these metrics further. The pixel locations in Fig. 5 that by eye lacked activity sub-BF (pixels 51, 53, 55, 60, 76) have *GR1* values ranging from 0.7 to 1.3. Considering the data more fully, the 0.7 (px 76) and 1.3 (px 55) values are different from 1 due to local dips in the data occurring at the frequency corresponding to BF/2. *GR2(dB)* was negative for both px 76 and px 55, and these points are considered to be in the frame, lacking sub-BF activity. *GR1* < 1 corresponds to expansive nonlinearity, and was attributable to local fluctuations. The red and blue color codes in the figures indicate sub-BF active (red) and sub-BF not-active (blue). In the remaining figures the two metrics are noted in the captions and/or in Table 1.

Fig. 6 is another measurement made through the RW with the optical axis nearly transverse. It reinforces observations of Fig. 5: sub-BF linearity in the BM and RL regions Fig. 5A&C, sub-BF nonlinearity in the OHC\DC region (Fig.5B) along with an amplitude trough and phase shift at high SPL (80 dB SPL). The phase differences shown in Fig. 6D&E are like those of Fig. 5, with nearly half cycle difference between the OHC\DC region and the RL from 30 to 50 kHz, the RL leading the BM by up to 0.2 cycles, and the 80 dB OHC\DC-region phase lifting to join the BM and RL phase at frequencies above those of the trough.

**Fig. 6.**
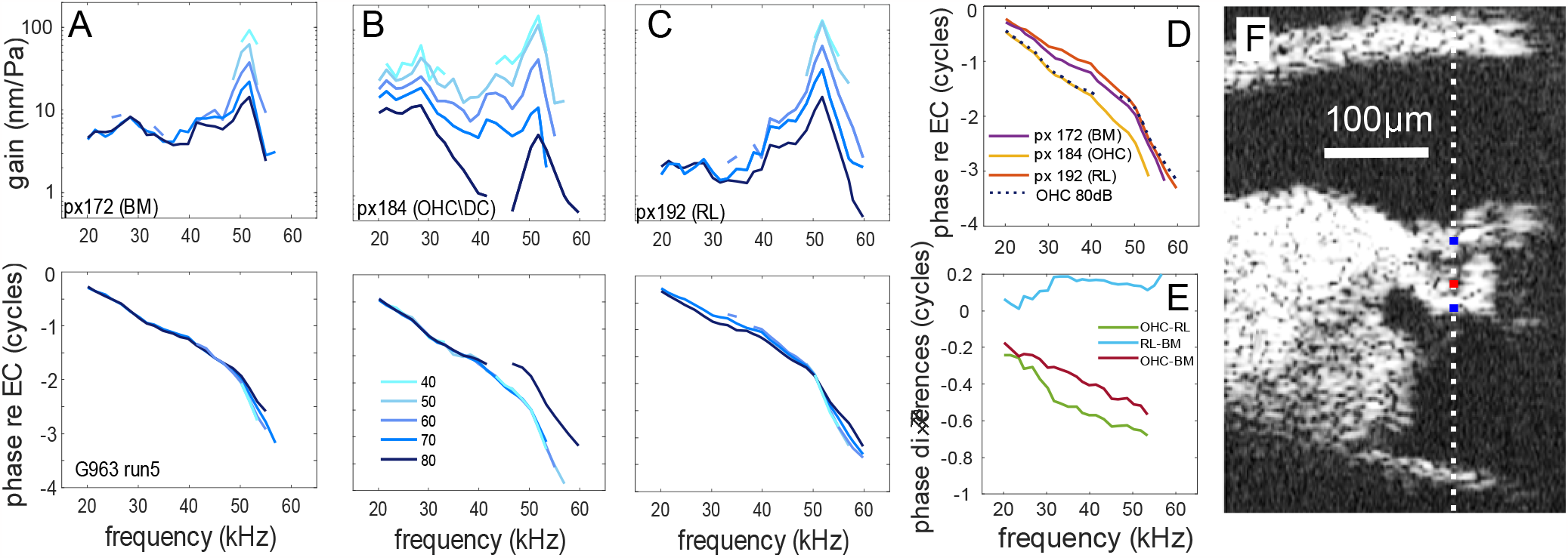
Another example of transverse data. A-C are gain and phase responses at the three pixels indicated in the B-scan in F, with the blue and red dots indicating locations without and with sub-BF nonlinearity respectively. D and E show phase comparisons. The multitone stimuli comprised 20 frequency components, spanning 20 to 60 kHz. Gain metrics: px172 GR1=1, GR2(dB)=-10; px184 GR1=1.5, GR2(dB)=4, px192 GR1=0.9, GR2(dB)=-24. Gerbil 963 run 5. BF= 52 kHz.

Figs. 5 and 6 show one of the primary observations of this study: when measured in the transverse direction, RL moves much like the BM. The RL and BM motion lacked the sub-BF nonlinearity and elevated gain of the OHC\DC region, lacked the high SPL amplitude trough, and compared to the large phase lag of the OHC\DC region, BM and RL motion were nearly in phase. However, RL motion differed from the BM: it was generally larger than BM motion in the BF peak, and smaller than BM motion sub-BF. The remaining plots will reinforce the primary observation and expand the observation to note the similarity of the quality of motion between BM and many regions of the OCC, which is the basis for the term “frame”.

Fig. 7A is another preparation measured through the RW with the optical axis nearly transverse. In this preparation a radial mapping of transverse motion was performed. Slices and pixel values are indicated and their locations noted in the B-scan (Fig. 7B). Three points showed clear sub-BF nonlinearity, they are slice 7 px 93 and 96 and slice 8 px 95, all in the OHC\DC region of the B-scan. Fig. 7C shows that these pixels contain the phase offsets and phase lift at 80 dB SPL to join the BM-like phase, as in Figs. 5&6. The responses at other points were generally similar to each other in gain shape and phase, and by the metrics were frame-like, but with some outliers. For example, slice 9 px107 shows sub-BF nonlinearity, but does not show the high SPL amplitude trough of the OHC\DC regions points, and the metrics of this point (*GR1=2*.*8, GR2(dB)=-8*) are ambiguous. Although it lacked the 80 dB phase lift, at other SPLs the phase of slice 9 p × 107 is like that of the OHC\DC points, so overall this pixel would be considered more hotspot than frame and is color-coded pink. From the B-scan, the location of slice 9 p × 107 is just below the outer tunnel within the tectal cells that join the lateral OCC to the RL, and its motion might be heavily influenced by fluid motion. Adjacent points within the RL (slice 6 p × 103 and slice 7 p × 104) are clearly frame-like by the metrics, lacking sub-BF nonlinearity, and with phases close to the BM phase (Fig. 7C). Slice 6 p × 94 is another unusual point, lacking sub-BF nonlinearity and frame-like by the metrics, but with relatively high gain values. In the B-scan this point appears to be nearly within the fluid space of the tunnel of Corti. From slice 9 through slice 15, a region spanning 60 μm laterally, all points except the unusual slice 9 p × 107 were frame-like: lacking sub-BF nonlinearity, with negative *GR2(dB)* values, and with phases close to those of the BM. Several of these points are deep within the OCC.

**Figure 7.**
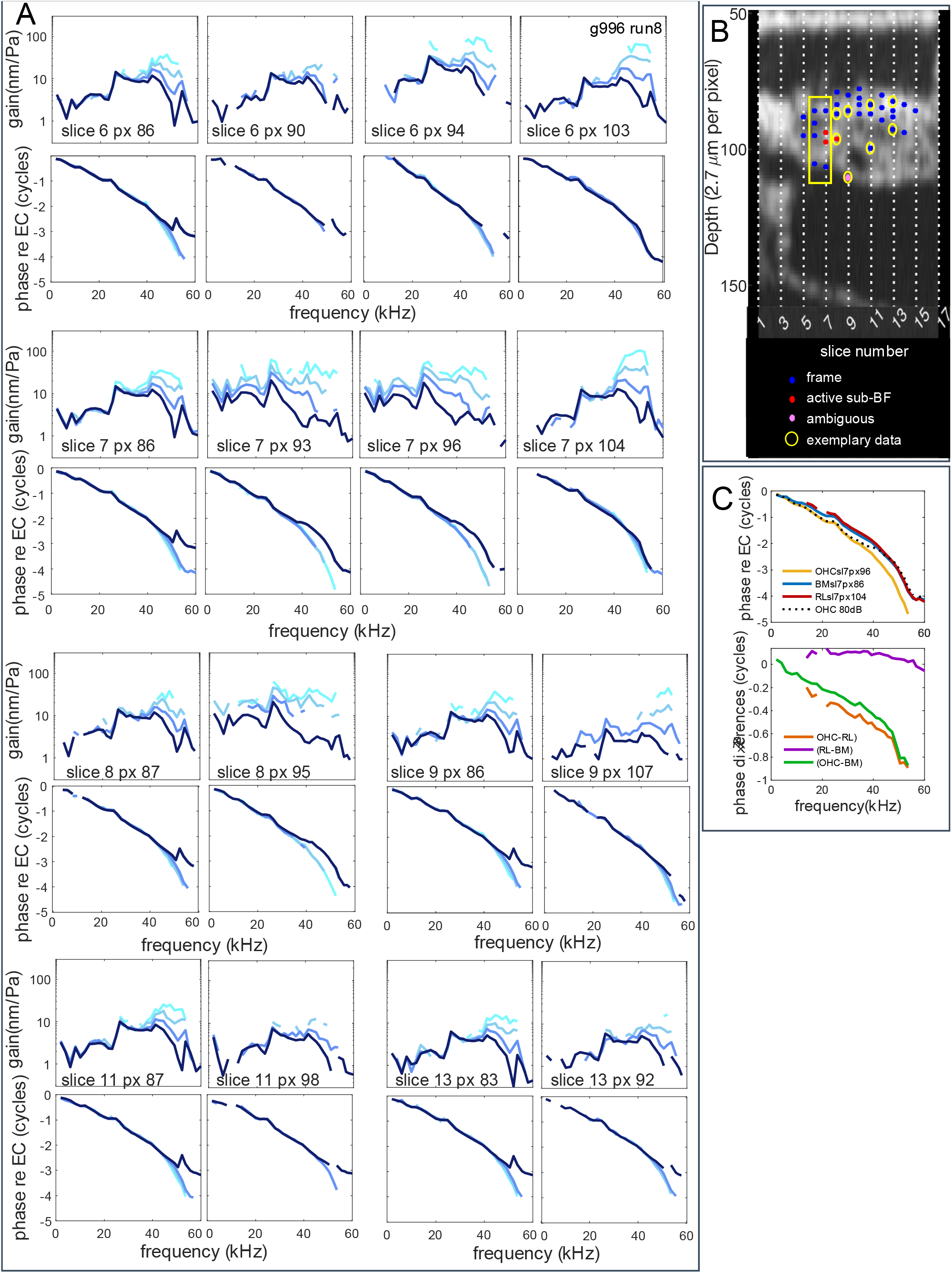
Frame map. A. Responses at locations indicated in the B-scan in (B). Multitone stimuli were applied at 50, 60, 70, 80 dB SPL (lighter to darker blue). In (A) the top panels show gain (displacement/EC pressure), bottom panels show phase relative to ear canal (EC) pressure. The lines in the B-scan are separated by 20 μm. The blue and red dots indicate regions without (blue) and with (red) sub-BF nonlinearity\activity. The pink dot is an ambiguous location. The yellow rectangle and circles identify locations where exemplary data are shown. C. Shows phases and phase differences from three locations in slice 7 (px 86, 96 and 104) representing BM, OHC\DC and RL. (70 dB SPL data were used for px86 and px104, 60 dB SPL for px96 to include a longer frequency range.) The multitone stimuli comprised 35 frequency components, spanning 2 to 60 kHz. Longitudinal, radial and transverse components were −0.31, −0.13, 0.94. Gerbil 996 run 8. Gain metrics are in Table 1.

In Fig. 8 we show two data sets from the ~ 23 kHz BF location from the same cochlea. Fig. 8 A-C data were taken through a cochleostomy above the stapedial artery (Fig. 2B), affording a nearly transverse view but limited to the very medial part of the OCC. These results did not include any locations with sub-BF nonlinearity; thus the three positions are within the OCC frame. Locations A and B are close to or within the BM, location C, 65 μm deeper than point A, is within or close to the RL, likely above the inner hair cell. The phases were similar across these locations.

**Figure 8.**
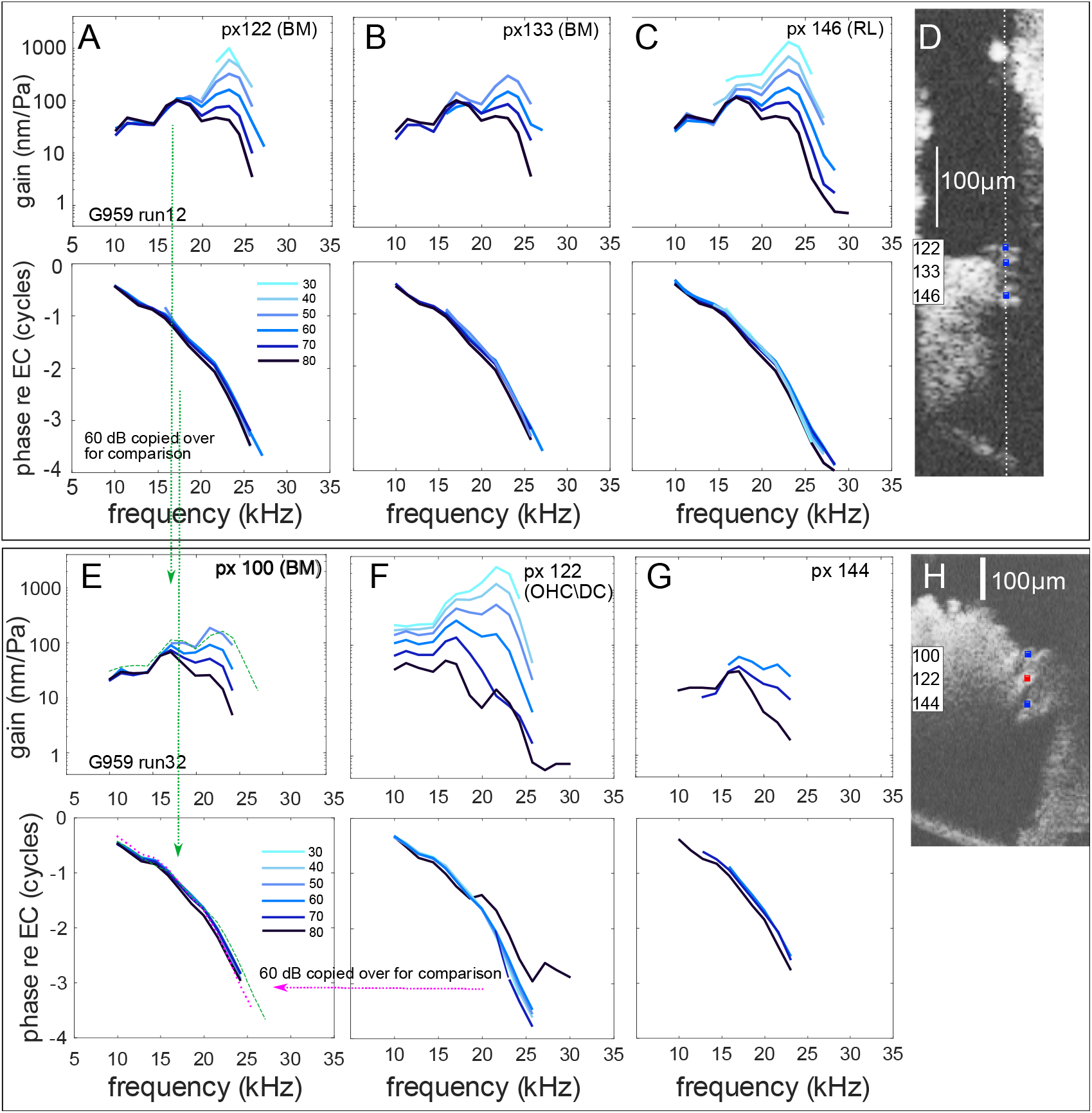
A-C. Motion responses taken through a cochleostomy as shown in Fig. 2B. D. Associated B-scan. Viewing axis was nearly transverse but not quantified. Results from three positions along one A-scan are shown in (A), (B) and (C) from locations indicated in the squares in the B-scan of (D). E-G. Responses at the same BF location as A-D, with data taken through the RW opening, with a substantial longitudinal component to the optical axis (not quantified). H. Associated B-scan. Results from three positions along one A-scan are shown in (E), (F) and (G) from locations indicated in the squares in the B-scan of (H). Anatomically, pixels 100, 122 and 144 are within the BM, OHC\DC region and lateral region respectively. In the BM panels (E), two comparisons are included. The pink dotted line is the phase responses from the OHC\DC region. The green dashed lines are the gain and phase responses copied from BM pixel 122 in (A). The multitone stimuli comprised 15 frequency components, spanning 10 to 30 kHz. Optical axis components were not measured. Metrics: run12 px 122 GR1=0.8, GR2(dB)=-6; run12 px 133 GR1=0.8, GR2(dB)=-8; run12 px 146 GR1=0.9, GR2(dB)=-6. run32 px100 GR1= 0.9, GR2(dB)= −5; run 32 px122 GR1= 1.6, GR2(dB)= 16; run32 px144 lacked a BF/2 point (~11 kHz), so is not included in the table of metrics, but when the 13 kHz 70 dB point is used as BF/2, GR1 < 1 and GR2(dB) is negative, which places this location in the frame, color-coded blue. Gerbil 959 run 12&32.

### 2. Optical axis containing substantial longitudinal component

Fig. 8 E-G data were from the same cochlea as Fig. 8 A-C, taken through the RW with the optical axis with a substantial longitudinal component, pointing apically to reach the same ~ 23 kHz BF location. The BM responses in Fig. 8E were like the BM responses measured through the cochleostomy, with a direct comparison of Fig. 8A 60 dB data included in Fig. 8E (green dashed lines). The BM is expected to move purely transversely, which would reduce the magnitude of the measured BM motion in Fig. 8E compared to Fig. 8A by the ratio of the transverse components of the optical axis in the two measurements (through RW/through cochleostomy). For example, if that value were 0.5 and these measurements were made at the same radial location across the BM, we would expect the green line to lie at twice the value of the 60 dB blue line. However, the Fig. 8A data were collected medial, closer to a rigid boundary, at a position that would move less than a more central radial position. Thus, there are two competing effects, and the result, with the green line from Fig. 8A 60 dB line slightly above the Fig. 8E 60 dB line, is reasonable. The Fig. 8 A-C data peaked at a just slightly higher BF than Fig. 8 E-G, due to inexact matching of BF location, but the overall similarity supports the view that the BM moves transversely, with motion that is similar, simply scaled, at different viewing angles (Frost et al., 2022; Cooper et al., 2018).

The B-scan in Fig. 8H is distorted from the transverse-radial view (Fig. 2A) due to the longitudinal\transverse optical axis, and the most obvious landmarks are the dark regions that correspond to the outer tunnel (below red dot) and tunnel of Corti (above-left of red dot), which sandwich the OHC\DC region (red dot). The motion at the BM (Fig. 8E) and ~120 μm within the OCC, in the region lateral to the outer tunnel (Fig. 8G) scaled linearly sub-BF, and are thus within the OCC frame. The OHC\DC-region motion (Fig. 8F) showed sub-BF nonlinearity. The OHC\DC region also showed an amplitude notch and phase shift at the highest SPL, but the notch and phase shift are different from those in Figs. 5-7; in Figs. 5-7 the 80 dB OHC phase shifted to join the BM phase, in Fig. 8F the OHC\DC-region phase was generally quite similar to BM phase (60 dB dotted pink line gives a direct comparison) and the notch-related phase shift at 80 dB shifted the phase away from the BM phase. The amplitude notch and phase shift in Fig. 8F are likely related to the OHC\DC-region motion being nearly perpendicular to the optical axis when the optical axis is substantially longitudinal (Frost et al., 2022, 2023a, 2023b). At frequencies below 15 kHz the OHC\DC region phase led the BM slightly, as in results from our previous reports with the longitudinal optical axis (Strimbu et al., 2020).

Fig. 9 is another example of data taken with the optical axis pointing apically down the cochlea to reach the ~ 25 kHz location. The first A-scan spanned the fluid regions that flank the OHCs (left line in Fig. 9H) and motion showed sub-BF nonlinearity in the OHC\DC region (Fig. 9B), but a deeper location seemingly close to or within the RL (Fig. 9C) was frame-like, without sub-BF nonlinearity. Locations probed in a second A-scan ~ 100 μm lateral, were all frame-like (Fig. 9 D-G). It was often not possible to visualize and thus probe the RL with the longitudinal approach and we include this data set to support the results from the more transverse view in which the RL does not present sub-BF nonlinearity and thus is within the OCC frame.

**Figure 9.**
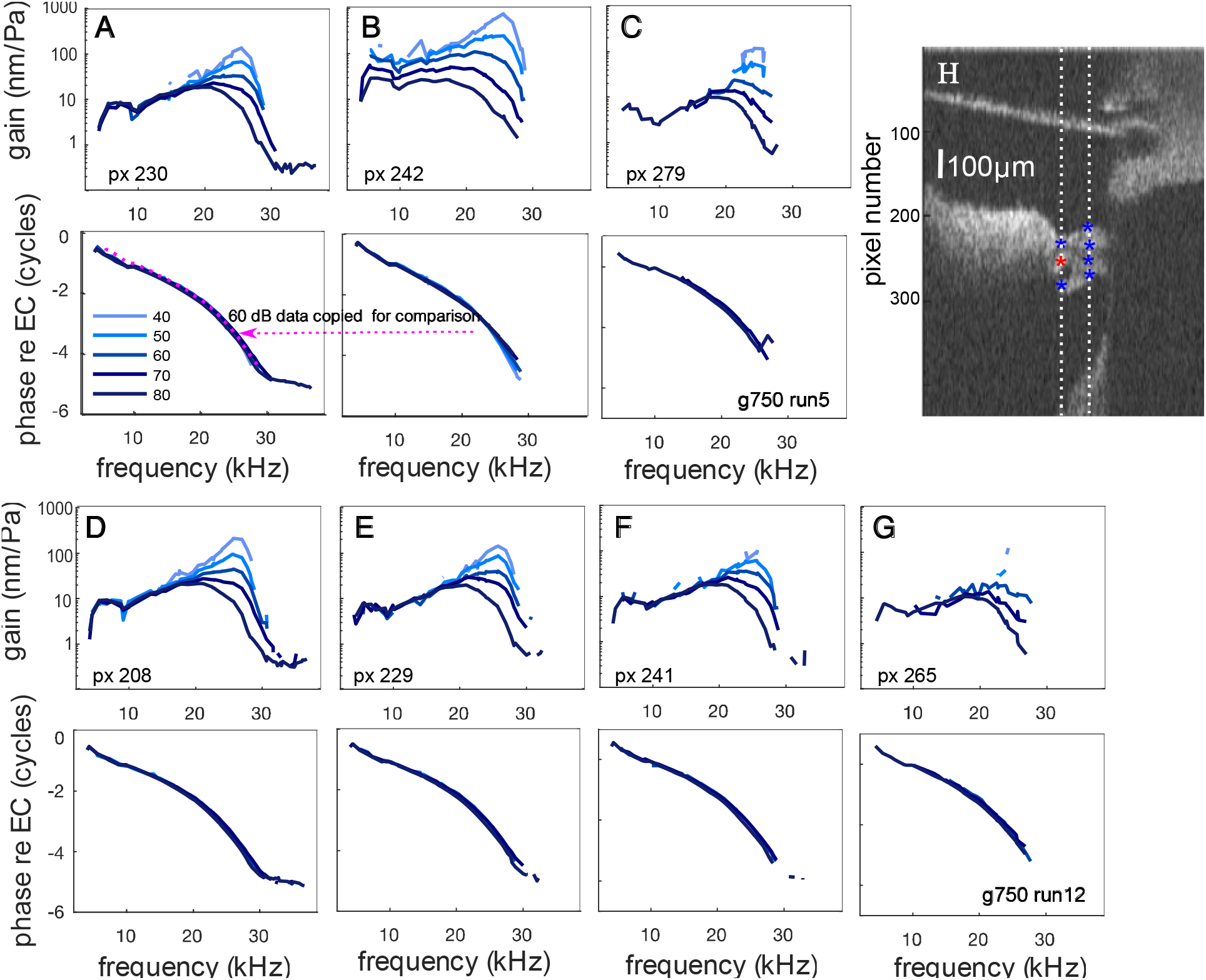
Another example in which data were taken through the RW opening, with a substantial longitudinal component to the optical axis. Responses were measured over two A-scans, indicated in the asterisks in the B-scan in H. A-C are from the left A-scan, D-G are from the right A-scan. The blue/red asterisks absence/presence of sub-BF nonlinearity (frame/hotspot). Gerbil 750, runs5&12. Metrics: run5 px230 GR1=0.9, GR2(dB)=-3; run5 px242 GR1=1.7, GR2(dB)=10; run5 px279 GR1=0.8, GR2(dB)=1. run12 px208 GR1= 0.9, GR2(dB)= 0; run 12 px229 GR1= 1.0, GR2(dB)= −6; run12 px241 GR1= 0.9, GR2(dB)= −4; run12 px265 GR1= 0.6, GR2(dB)= 3.

The measurements in Fig. 10 were made with the optical axis substantially longitudinal to reach the ~ 31 kHz BF location. Motion gain data are shown in the plots to the left of the orienting B-scan. Figs. 8 and 9 showed that phase in the more longitudinal approach is less revealing than in the transverse approach, thus in Fig. 10 phase data were not included. (All locations in Fig. 10 contained traveling wave phase accumulation.) The dark fluid-filled spaces in the B-scan that flank the OHC\DC region were used to identify that region. Three exemplary data sets are shown that possessed clear sub-BF nonlinearity (red points with yellow circles, slice 4 p×189, slice 6 p×179, slice 7 p×172). Slice 4 p×191 is close to or within the RL. By eye this point was ambiguous in terms of sub-BF nonlinearity, and the metrics were also ambiguous, but lean toward a “frame” designation (GR1=1.3 (borderline), GR2(dB)=-2 (in frame)). This point was color-coded pink. Slice 3 p×197 was also deemed ambiguous, by metrics and by eye. These ambiguous points might be at or close to the RL, but the longitudinal view obscures the familiar cross-sectional anatomy too much to know. From slice 8 through slice 12, all points were within the frame, and several of these points are deep within the OCC.

**Figure 10.**
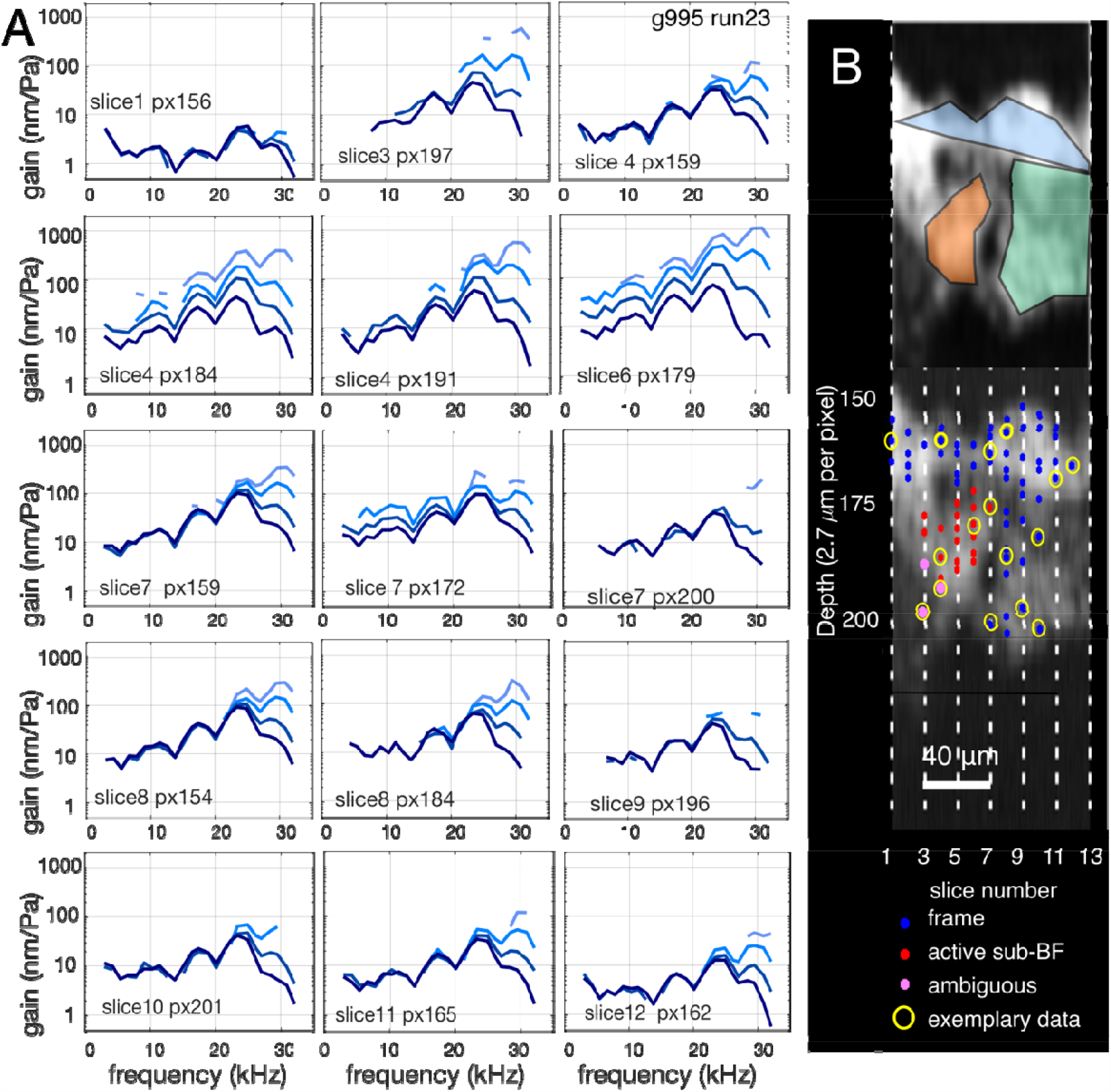
Frame map. A. Gain responses at locations indicated in the B-scan in (B). Multitone stimuli were applied a 40, 53, 67, 80 dB SPL (lighter to darker blue. The blue and red dots indicate regions without (blue) and with (red) sub-BF nonlinearity\activity. The pink dots are ambiguous locations. The yellow circles identify locations where exemplary data are shown. Optical axis longitudinal, radial and transverse components were 0.85, 0.24, 0.47. Gerbil 995 run 23. Gain metrics are in Table 1.

## Discussion

### A. OHC-region behavior when optical axis is relatively transverse -- complex differences to find internal motion

The OHC\DC-region measurements made along a relatively transverse optical axis (Figs. 5-7) possessed magnitude troughs and associated phase shifts. These characteristics suggest that the measured OHC\DC-region motion can be considered as a sum of an internal OHC (hotspot) motion and the motion of the frame. The BM is the primary structural element of the OCC frame and is taken as the reference frame structure. Based on its anatomy, with its set of tightly packed and radially-oriented collagen fibers (de Sousa Lobo Querido et al., 2023; Dreiling et al., 2002), the BM is expected to move primarily in the transverse direction, and this was supported by the BM results in Fig. 8. Taking the complex difference, OHC\DC-region minus BM motion, provides an estimate of the internal transverse motion of the OHC\DC-region. This analysis is illustrated in the cartoon of Fig. 11, and the analysis was done for the data of Figs. 5-7 and shown in Fig. 12. In Figs. 11&12, BM motion is blue, OHC\DC-region motion is red, and their complex difference is gold. Because the BM and OHC\DC-region motions can be nearly out of phase, the magnitude of their difference was often larger than the magnitudes of the OHC\DC region and BM motion -- in particular, the BF peak in the gold curves in Fig. 12A at 50-60 dB SPL, in Fig. 12B at 50-70 dB SPL and Fig. 12C at 50-60 dB SPL. The (red) OHC\DC magnitude trough is either not present or reduced in the calculated internal motion (gold), and the phase shift observed in the measured OHC\DC motion is not present in the internal motion. In the 80 dB curves of Fig. 12A-C the reason for the trough in measured OHC\DC-region motion (red) is easily traced to the equal amplitudes and half cycle phase differences of the blue and gold curves of BM and internal OHC\DC-region motion. Thus, the results of the complex difference analysis clarified oddities of the measured motion.

**Figure 11.**
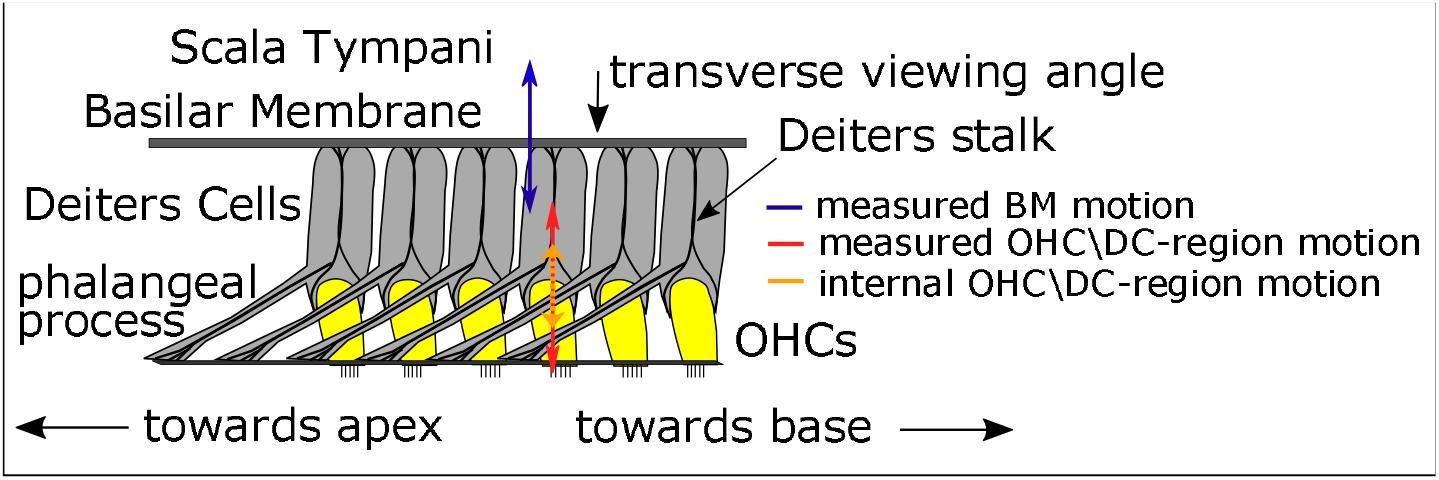
Longitudinal-transverse cartoon of the OCC from the BM to the RL.

**Figure 12.**
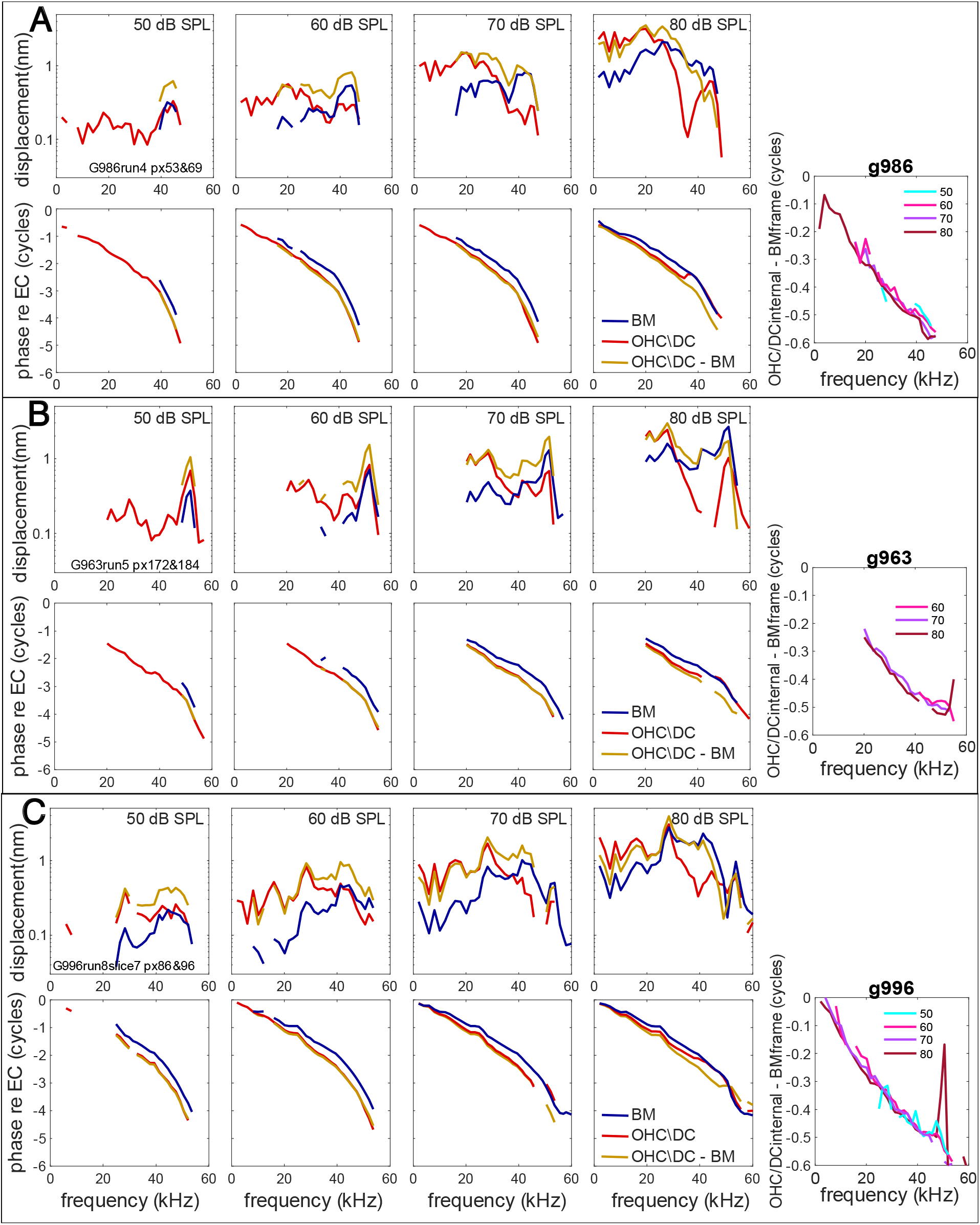
Complex difference between OHC\DC region and BM responses was taken to reveal internal OHC\DC motion. In each preparation the responses going into the difference were measured along a single A-scan as in the motion data of Fig.5 (Gerbil 986), Fig.6 (Gerbil 963) and Fig.7 (Gerbil 996). Results are shown in panel sets A,B,C, respectively. Each panel set has subpanels corresponding to different SPL and within each subpanel, the OHC\DC and BM data are shown in red and blue respectively, and the OHC-BM complex difference is shown in gold. To the far right the phase of the internal OHC\DC motion (hotspot) is plotted referenced to the BM (frame).

At the far right of the panel sets is the phase of the internal OHC\DC (hotspot) motion relative to the BM (frame). This phase is similar across sound levels (50-80 dB SPL) and preparations. The phase is close to zero at the lowest frequencies and decreases generally smoothly and steadily to a value of −0.5 cycles just below BF and continues decreasing to a value of ~ −0.6 cycles. The significance of this hotspot motion, moving largely in the opposite direction relative to the motion of the frame, is not known. It is possible that it is simply a byproduct, an epiphenomenon of the active process. Conversely, fluid moving to accommodate the hotspot motion may be critical to the longitudinal flow of energy towards the best place. Several recent cochlear models have explored the role of active fluid flow in frequency tuning (Altoe et al., 2022; Guinan, 2022; He et al., 2022). The findings here add to the experimental data guiding the models.

The analysis of Figs. 11&12 could only be applied to the data sets taken with a relatively transverse view. For the data sets taken with a significantly longitudinal optical axis (Figs. 8 - 10), an analysis to reconstruct transverse and longitudinal motion has been devised but requires more extensive data to implement (Frost et al., 2023a,b).

### B. Comparison with previous results

The measurements of Figs. 8 - 10 were made with an approach through the RW opening and angling longitudinally to reach the ~ 25 kHz location. This approach was used in several of our previous studies and the results presented here are similar to those and the earlier results of Cooper et al., (2018), who also used multitone stimuli. Because of the steep longitudinal optical angle, the RL is often not observable in this approach, and in our previous reports we emphasized OHC\DC and BM regions.

The measurements of Figs. 5-7 were made with a nearly transverse approach through the RW. This approach affords a transverse-radial B-scan, and the RL region can be identified. This approach is like that of Cho and Puria, 2022, and the findings were similar. Cho and Puria used single-tone stimuli and in their case the presence of sub-BF activity in the OHC\DC region was observed via *post-mortem* reduction in sub-BF motion. In the RL and BM, sub-BF motion was not reduced post-mortem. The Cho and Puria data therefore are consistent with our RL data, and support the statement that the RL does not possess sub-BF activity, and is thus within the OCC frame.

The Fig. 8A-C observations were made through a cochleostomy, to make transverse measurements at the ~ 25 kHz location. The view was limited to the medial OCC and did not include the OHC region. Measurements were made at the BM and 65 μm deeper within the OCC, likely at the RL adjacent to the pillar cells. Sub-BF responses were linear; all positions were within the frame. He et al., 2022 made measurements at this BF location through a cochleostomy after surgically displacing the stapedial artery to afford a more complete view of the OCC. Their system did not include B-scan imaging, and the RL and BM were identified as reflective surfaces with a reasonable separation distance. The region they identified as the RL exhibited sub-BF nonlinearity from 70 to 80 dB SPL, and at BF/2, the RL responses were elevated by ~ 16 dB relative to BM. Previous work by He et al. (2018) had shown a ~ 20 *dB post-mortem* reduction in sub-BF RL motion. Thus, responses in the region He et al. identified as RL displayed sub-BF activity. This finding conflicts with our observations that the RL exhibits little or no sub-BF activity. It is notable that in our Fig. 5, along a single A-scan, p × 74 with sub-BF activity was only 6 μm from p × 76 lacking sub-BF activity. Thus, a B-scan and ability to measure from many axial locations simultaneously, as is afforded with spectral domain OCT, is an advantage. Regardless, this discrepancy bears further study.

### C. Significance of the frame for hair cell excitation

In the base of the cochlea, neural tuning is similar to BM motion tuning, especially in the BF peak (Naryanan et al., 1998). Hair cells are stimulated by the pivoting motion of their stereocilia, which in Fig. 2A stand below the RL and insert (OHCs) or nearly insert (IHCs) into the TM (Lim, 1972). In the results of this study, the RL lacked sub-BF activity. Thus, it seems that hair cell stereocilia, both IHC and OHC, are not stimulated by the relatively large and nonlinear sub-BF motion that is present in the OHC\DC region. The sub-BF nonlinearity observed in local cochlear microphonic (Fallah et al., 2019) would in that case be due solely to MET nonlinearity, and not to nonlinear mechanical input to the OHC stereocilia. In damaged cochleae high-frequency auditory neurons can become hyper-sensitive to sub-BF tones and contain temporal firing patterns including phase locking that are not present in healthy high-frequency auditory neurons (Versnel et al., 1996; Kale and Heinz 2010). It is possible that in damaged cochleae the frame can become compromised, allowing OHC activity to overstimulate stereocilia at sub-BF frequencies, leading to pathological auditory nerve responses. On the other hand, a lack of sub-BF activity in stereocilia stimulation is at odds with studies finding sub-BF responses of high frequency auditory nerve fibers could be inhibited by medial olivocochlear activity and suppressed by a low frequency suppressor (Nam and Guinan, 2018).

The measurements reported here did not include the TM, which was not reflective enough to provide the vibration responses necessary to evaluate sub-BF nonlinearity. However, in reports from the mouse 9 kHz BF region, with an optical axis containing radial and transverse components, the TM vibrated like the BM (like the frame) and not like OHC region, in terms of nonlinearity and *post-mortem responses* (Dewey et al., 2018; Dewey, 2022). This supports the expectation that the TM is within the frame and suggests that the frame concept is applicable to the 9 kHz region of the mouse cochlea as well as the cochlear base of gerbil and guinea pig. However, earlier results in mouse from the same group did observe sub-BF nonlinearity in the TM and RL (Lee et al., 2016). Overall, OCT measurements are still novel, the 3-dimensional motion in the OHC\TM\RL region appears to be spatially rapidly changing, and the behavior of the motion in that region is still in the process of being ironed out. Nevertheless, the findings of this study, and the study of Cho and Puria (2022), indicate that in the base of the gerbil cochlea the RL is shielded from the sub-BF activity of the OHC\DC region, and in the terminology forwarded here, the RL is part of the OCC frame.

### D. Physical ramifications of OCC frame

Nonlinear motion abutting tissue exerts nonlinear forces on that tissue. In Fig. 11, the internal nonlinear OHC motion will transmit nonlinear forces to the Deiters cells and RL. However, sub-BF nonlinear motion is not observed at the BM or RL or other components of the OCC frame. Thus, sub-BF forces on frame structures must be dominated by passive forces. A primary force on the BM is the pressure on its scala tympani surface. This pressure has been measured in active cochleae and is tuned and nonlinear in the BF peak but sub-BF is linear and passive (*unchanging post-mortem*) (Olson, 2001). The slow wave pressure on the scala media side of the OCC is theoretically equal in magnitude to the pressure in scala tympani, and opposite in phase. Pressure measured in scala media was consistent with that expectation, although the preparation was in an unavoidably damaged condition (Kale and Olson, 2015). Pressure and BM motion are partners in the cochlear traveling wave, and previous studies have reported that sub-BF activity is not transported by the wave (Fallah et al., 2019; Dewey et al., 2018, Guinan, 2020). Thus, it is reasonable that at sub-BF frequencies the pressure, which is not active sub-BF, is a dominant force on the OCC.

The pillar cells and the Deiters stalk, composed of structural proteins (microtubules, actin and intermediate filaments) anchor the RL to the BM (Engstrom and Wersall, 1958; Parsa et al., 2012; Zhou et al., 2022). The physical properties and orientation of these structures might further constrain the RL to move more like the BM, and not like the OHCs. The phalangeal processes of the DCs course apically (Fig. 11) and as the cochlear wave travels apically, their mechanical effect will be to impose basal BM motion on the more apical RL location. Thus, in transverse measurements of the RL and BM vibration, the RL would be predicted to lead the BM. The value of the predicted phase lead depends on the traveling wave wavelength, which is ~ 0.4 mm at frequencies getting close to the BF (Ren 2002) and longer at lower frequencies. The phalangeal processes in the gerbil base span a longitudinal distance of ~ .08 mm (Wang et al., 2016). Thus, at frequencies approaching the BF the phalangeal processes span ~ 0.2 wavelength, corresponding to 0.2 cycle. This is close to the measured RL re BM phase lead in Figs. 5E, 6E and 7C. This speculative prediction is dimmed by the fact that in Cho and Puria (2022) the phase lead increased at low frequencies, rather than approaching zero as the thinking above would have it. They also found the phase lead disappeared *post-mortem*, which would not be the case if the intracellular anchoring structures were unchanged *post-mortem*. Nevertheless, a possible role for the Deiters stalks in imposing BM-like motion on the RL is worthy of further study. The pillar cells most certainly have such a role based on their significant stiffness (Olson and Mountain, 1994) and supported by recent across-RL measurements (Cho and Puria, 2022).

A central question of cochlear mechanics remains: how does the BM, and OCC frame in general, respond to OHC active forces only at frequencies in the BF peak, when OHC active forces are present through the full range of sub-BF frequencies? Mathematical models and concepts can replicate this phenomenon (Nankali et al., 2020; Yoon et al., 2011, deBoer and Nuttall, 2000; Altoe et al., 2022; Sisto et al., 2021; Dong and Olson, 2013) but their physiological underpinnings remain tenuous. OCT has delivered a wealth of new data regarding the coordinated responses of OHCs, supporting cells and their surrounding fluid and extracellular structures. In the current analysis, OCT observations have advanced the concept of separate “internal-hotspot” and “frame” motions, whose separation is likely essential for healthy cochlear function.

## Acknowledgments

Funding was provided by the NIDCD grants R01 DC015363 (Elizabeth Olson PI) and F31 DC020621 (Brian Frost PI). Funding was provided by the Emil Capita Foundation.

## References

Altoe A, Dewey JB, Charaziak KK, Oghalai JS & Shera CA (2022) Overturning the mechanisms of cochlear amplification via area deformations of the organ of Corti. J Acoust Soc Am 152: 2227–2239.

Cho NH & Puria S (2022) Cochlear motion across the reticular lamina implies that it is not a stiff plate. Sci Rep 12: 18715.

Cooper NP Vavakou A & van der Heijden M (2018) Vibration hotspots reveal longitudinal funneling of sound-evoked motion in the mammalian cochlea Nature communications 9(1) 1–12

De Boer E & Nuttall AL (2000) The mechanical waveform of the basilar membrane III Intensity effects. J Acoust Soc Am 107:1497–1507.

de Sousa Lobo Querido R, Ji X, Lakha R, Goodyear RJ, Richardson GP, Vizcarra CL & Olson ES (2023) Visualizing collagen fibrils in the cochlea’s tectorial and basilar membranes using a fluorescently labeled collagen-binding protein fragment. JARO 24: 1–17.

Dewey JB, Xia A, Muller U, Belyantseva IA, Applegate BE & Oghalai JS (2018) Mammalian auditory hair cell bundle stiffness affects frequency tuning by increasing coupling along the Length of the cochlea. Cell Reports 23, 2915–2927.

Dewey JB, Applegate BE & Oghalai JS (2019) Amplification and suppression of traveling waves along the mouse organ of corti: evidence for spatial variation in the longitudinal coupling of outer hair cell-generated forces. J Neurosci 39: 1805–1816.

Dewey JB (2022) Cubic and quadratic distortion products in vibrations of the mouse cochlear apex. JASA Express Letters 2(11) 114402,1-7.

Dong W & Olson ES (2008) Supporting evidence for reverse cochlear traveling waves. J Acoust Soc Am 123: 222–240.

Dong W & Olson ES (2013) Detection of cochlear amplification and its activation. Biophysical J 105: 1067–1078.

Dong W & Olson ES (2016) Two-tone suppression of simultaneous electrical and mechanical responses in the cochlea. Biophysical J 111: 1805–1815.

Dreiling FJ, Henson MM & Henson OW Jr (2002) The presence and arrangement of type II collagen in the basilar membrane. Hearing Research 166:166–180.

Engstrom H & Wersall J (1958) The ultrastructural organization of the organ of Corti and of the vestibular sensory epithelia. Experimental Cell Research 14:460–492.

Fallah E, Strimbu CE & Olson ES (2019) Nonlinearity and amplification in cochlear responses to single and multi-tone stimuli. Hearing Research 377:271–281.

Fallah E, Strimbu CE & Olson ES (2021) Nonlinearity of intracochlear motion and local cochlear microphonic: Comparison between guinea pig and gerbil. Hearing Research 405: 108234.

Fettiplace R & Kim KX (2014) The physiology of mechanoelectrical transduction channels in hearing. Physiological Reviews 94: 951–986.

Frank G, Hemmert W & Gummer AW (1999) Limiting dynamics of high-frequency electromechanical transduction of outer hair cells. PNAS 96: 4420–4425.

Frost BL, Strimbu CE & Olson ES (2022) Using volumetric optical coherence tomography to achieve spatially resolved organ of Corti vibration measurements. J Acoust Soc Am 151: 1115–1124.

Frost BL, Strimbu CE & Olson ES (2023a) Reconstruction of transverse-longitudinal vibrations in the organ of Corti complex via optical coherence tomography. J Acoust Soc Am 153: 1347–1360.

Frost BL, Strimbu CE & Olson ES (2023b) Erratum: Reconstruction of transverse-longitudinal vibrations in the organ of Corti complex via optical coherence tomography [J Acoust Soc Am 153 1347-1360]. J Acoust Soc Am 153: 2537.

Guinan JJ Jr. (2022) Cochlear amplification in the short-wave region by outer hair cells changing organ-of-Corti area to amplify the fluid traveling wave. Hearing Research 426: 108641.

Hallworth R, Evans BN and Dallos P (1993) The location and mechanism of electromotility in guinea pig outer hair cells. J Neurophysiology 70: 549–558.

He W, Kemp D & Ren T (2018) Timing of the reticular lamina and basilar membrane vibration in living gerbil cochleae. eLife 7:e37625.

He W, Burwood G, Porsov EV, Fridberger A, Nuttall AL & Ren T (2022) The reticular lamina and basilar membrane vibrations in the transverse direction in the basal turn of the living gerbil cochlea. Sci Rep 12: 19810.

He W, Burwood G, Fridberger A, Nuttall AF and Ren T (2022) An outer hair cell-powered global hydromechanical mechanism for cochlear amplification. Hear Res 432: 10847 pp 1-11.

Iwasa KH & Adachi M (1997) Force generation in the outer hair cell of the cochlea. Biophysical J 73: 546–555.

Kale S & Heinz MG (2010) Envelope coding in auditory nerve fibers following noise-induced hearing loss. JARO 11: 657–673.

Kale S & Olson ES (2015) Intracochlear scala media pressure measurement: Implications for models of cochlear mechanics. Biophys J 109: 2678–2688.

Lee HY, Raphael PD, Xia A, Kim J, Grillet N, Applegate BE, Bowden AKE & Oghalai JS (2016) Twodimensional cochlear micromechanics measured in vivo demonstrate radial tuning with the mouse organ of Corti. J Neuro 36:8160–8173.

Lin NC, Hendon CP and Olson ES (2017) Signal competition in optical coherence tomography and its relevance for cochlear vibrometry. JASA 141: 395–405.

Nankali A, Wang Y, Strimbu CE, Olson ES & Grosh K (2020) A role for tectorial membrane mechanics in activating the cochlear amplifier. Sci Rep 10: 17620.

Nam H, Guinan JJ Jr (2018) Non-tip auditory-nerve responses that are suppressed by low-frequency bias tones originate from reticular lamina motion. Hearing Research 358: 1–9.

Narayan SS, Temchin AN, Recio A & Ruggero MA (1998) Frequency tuning of basilar membrane and auditory nerve fibers in the same cochleae. Science 28: 1882–1884.

Olson E S (2001) Intracochlear pressure measurements related to cochlear frequency tuning. J Acoust Soc Am 110(1) 349–367.

Olson ES and Mountain DC (1994) Mapping the cochlear partition’s stiffness to its cellular architecture. J Acoust Soc Am 95: 395–400.

Parsa A, Webster P & Kalinec F (2012) Deiters cells tread a narrow path: The Deiters cells-basilar membrane junction. Hearing Research 290: 13–20.

Rabbitt RD (2022) Analysis of outer hair cell electromechanics reveals power delivery at the upper frequency limits of hearing. J R Soc Interface 19: 20220139.

Rhode WS (2007) Basilar membrane mechanics in the 6–9 kHz region of sensitive chinchilla cochleae. J Acoust Soc Am 121:2792–2804.

Sisto R, Belardinelli D & Moleti A (2021) Fluid focusing and viscosity allow high gain and stability of the cochlear response. J Acoust Soc Am 150: 4283–4296.

Strimbu CE Wang Y & Olson ES (2020) Manipulation of the endocochlear potential reveals two distinct types of cochlear nonlinearity Biophysical J 119: 1–15.

van der Heijden M & Joris PX (2003) Cochlear phase and amplitude retrieved from the auditory nerve at arbitrary frequencies. J Neurosci 23(27) 9194–9198.

Strimbu CE & Olson E S (2022) Salicylate-induced changes in organ of Corti vibrations. Hearing Research 423: 108389.

van der Heijden M & Joris PX (2003) Cochlear phase and amplitude retrieved from the auditory nerve at arbitrary frequencies. J Neurosci 23:9194–9198.

Versnel H, Prijs VF & Schoonhoven R (1997) Auditory-nerve fiber responses to clicks in guinea pigs with a damaged cochlea J Acoust Soc Am 101: 993–1009.

Versteegh CP & van der Heijden M (2012) Basilar membrane responses to tones and tone complexes: nonlinear effects of stimulus intensity. JARO 13: 785–798.

Wang Y, Steele CR & Puria S (2016) Cochlear outer-hair-cell power generation and viscous fluid loss. Sci Rep 6:19475.

Warren RL, Ramamoorthy S, Ciganovic N & Lim DJ (1972) Fine morphology of the tectorial membrane: its relationship to the organ of Corti. Arch Otolaryngology 96:199–215.

Yoon YJ, Steele CR & Puria S (2011) Feed-forward and feed-backward amplification model from cochlear cytoarchitecture: an interspecies comparison. Biophysical J 100: 1–10.

Zhou W, Jabeen T, Sabha S, Becker J & Nam J-H (2022) Deiters cells act as mechanical equalizers for outer hair cells J Neurosci 42:8361–8372.

